# Disruption of folate metabolism causes poor alignment and spacing of mouse conceptuses for multiple generations

**DOI:** 10.1101/2021.06.11.448112

**Authors:** Amy L. Wilkinson, Katerina Menelaou, Joanna Rakoczy, Xiu S. Tan, Erica D. Watson

## Abstract

Abnormal uptake or metabolism of folate increases risk of human pregnancy complications, though the mechanism is unclear. Here, we explore how defective folate metabolism influences early development by analysing mice with an *Mtrr*^*gt*^ hypomorphic mutation. MTRR is necessary for methyl group utilisation from the folate cycle, and the *Mtrr*^*gt*^ allele disrupts this process. We show that the spectrum of phenotypes previously observed in *Mtrr*^*gt/gt*^ conceptuses at embryonic day (E) 10.5 is apparent from E8.5 including developmental delay, congenital malformations, and placental phenotypes (e.g., eccentric chorioallantoic attachment). Notably, we report misalignment of some *Mtrr*^*gt*^ conceptuses within their implantation sites from E6.5. The degree of skewed growth occurs across a continuum, with eccentric chorioallantoic attachment now re-characterised as a severe form of conceptus misalignment. Additionally, some *Mtrr*^*gt/gt*^ conceptuses display twinning. Therefore, we implicate folate metabolism in blastocyst orientation and spacing at implantation. Embryo development is influenced by skewed growth since developmental delay and heart malformations (but not neural tube defects) associate with severe misalignment of *Mtrr*^*gt/gt*^ conceptuses. Patterning of trophoblast lineage markers is largely unaffected in skewed *Mtrr*^*gt/gt*^ conceptuses at E8.5 indicating trophoblast differentiation was normal when misaligned. Typically, the uterus guides conceptus orientation. Accordingly, we manipulate the maternal *Mtrr* genotype and assess conceptus alignment. *Mtrr*^*+/gt*^, and *Mtrr*^*gt/gt*^ mothers, plus *Mtrr*^*+/+*^ mothers, exhibit misaligned conceptuses at E6.5. While progesterone and/or BMP2 signalling required for decidualisation might be disrupted, normal gross decidual morphology, patterning, and blood perfusion is evident regardless of conceptus alignment, arguing against a uterine defect. Given the important finding that *Mtrr*^*+/+*^ mothers also display conceptus misalignment, a grandparental effect is explored. Multigenerational phenotype inheritance is characteristic of the *Mtrr*^*gt*^ model, though the mechanism remains unclear. Genetic pedigree analysis reveals that severe skewing associates with the *Mtrr* genotype of either maternal grandparent. Moreover, misalignment is independent of the uterus and instead is attributed to an embryonic mechanism based on blastocyst transfer experiments. Overall, our data indicates that abnormal folate metabolism influences conceptus orientation over multiple generations with implications for subsequent development. Our study casts light on the complex role of folate metabolism during development beyond a direct maternal effect.

## INTRODUCTION

Maternal folate deficiency in humans is associated with increased risk of congenital malformations and pregnancy complications, such as placental abruption, haemorrhage, and pre-eclampsia (Wen et al., 2008). These complications potentially lead to increased risk of spontaneous abortion, miscarriage, or pre-term birth (Bukowski et al., 2009). Yet, how folate metabolism functions in pregnancy establishment is not well understood. Folate, a vitamin that underpins one-carbon metabolism, is required for thymidine synthesis (Stover, 2004) and transfer of one-carbon methyl groups for cellular methylation reactions (Ducker and Rabinowitz, 2017). Therefore, folate is required for rapid cell division and epigenetic regulation of gene expression, which are both important for feto-placental development.

At implantation, human and mouse blastocysts embed into the uterine wall to optimize fetal access to maternal resources. Mouse blastocysts attach to the antimesometrial uterine epithelium via the mural trophectoderm (opposes the inner cell mass (ICM)) and invade into the decidualising endometrium by embryonic day (E) 4.5 (Smith, 1985; Cross et al., 1994). In contrast, human blastocysts attach to the uterus via the polar trophectoderm (contacts the ICM) (Hertig et al., 1956). Mouse litters consist of 5-20 embryos, each form their own placenta on the mesometrial side of the uterus (Smith, 1985) and must be equally spaced within the uterus for ideal growth. Spacing mechanisms are not well understood but likely involve mechanical forces generated by the uterus and cross talk between blastocysts and the uterine epithelium (Flores et al., 2020). After implantation, the ICM will form the embryo proper, with the progenitor cells of the cardiovascular and nervous systems among the first to develop (McDole et al., 2018). Simultaneously, the polar trophectoderm forms the ectoplacental cone (EPC) and extra-embryonic ectoderm, which together consist of progenitor populations of the mature placenta (Watson and Cross, 2005). The EPC is a source of invasive trophoblast cells involved in uterine spiral artery remodelling and gives rise to the endocrine junctional zone cells of the mature placenta (Adamson et al., 2002; Simmons and Cross, 2005; Simmons et al., 2007). The extra-embryonic ectoderm gives rise to the chorion, which will form the labyrinth layer with the fetal vasculature after the allantois attaches to the chorion in a process called chorioallantoic attachment (Watson and Cross, 2005). The labyrinth is the site of nutrient and gas exchange between maternal and fetal blood circulations (Watson and Cross, 2005). The orientation of the conceptus is determined in a stereotypical manner aligning with the antimesometrial-mesometrial axis of the decidual swelling. The chorion located at the base of the EPC with the tip of the EPC extending upwards through the mesometrial decidua. The embryo proper is surrounded by the yolk sac below the chorion on the antimesometrial side of the implantation site. The extent to which conceptus alignment influences embryo and placenta development is not well understood.

Furthermore, whether folate metabolism plays a role in implantation, spacing or orientation remains unclear, as only a few models of folate deficiency have been explored in this context. During its metabolism, folate is converted to 5-methyltetrahydrofolate by methylene tetrahydrofolate reductase (MTHFR). Methionine synthase (MTR) then catalyses the transfer of the methyl group from 5-methyltetrahydrofolate to homocysteine to produce methionine (Shane and Stokstad, 1985). Methionine is the precursor of S-adenosylmethionine, the methyl donor of all methylation reactions (Ducker and Rabinowitz, 2017). Importantly, methionine synthase reductase (MTRR) ensures progression of both folate and methionine cycles by continued activation of MTR through reductive methylation of its vitamin B_12_ co-factor (Yamada et al., 2006). Mutations in human *MTHFR, MTR*, and *MTRR* genes are linked to pregnancy complications (Ren and Wang, 2006; Furness et al., 2008; Kim et al., 2013; Wang et al., 2015), yet the mechanism of one-carbon metabolism in this context requires further study.

Several mouse models of defective folate uptake, transport or metabolism suggest involvement of folate metabolism in early pregnancy. For instance, dietary folate deficiency in mice is associated with sub-fertility, increased embryonic lethality attributed to defects in decidualisation (Gao et al., 2012; Geng et al., 2015), and low weight placentas with shallow trophoblast invasion (Pickell et al., 2009). Mutations in mouse genes associated with folate uptake or metabolism display gross placental phenotypes (e.g., *Slc19a1*^*-/-*^, *Mthfr*^*-/-*^ and *Mtrr*^*gt/gt*^) or are yet-to-be assessed for defects beyond the embryo proper (e.g., *Folbp1*^*-/-*^, *Folbp2*^*-/-*^, *Slc46a1*^*-/-*^, and *Mtr*^*-/-*^) (Piedrahita et al., 1999; Swanson et al., 2001; Elmore et al., 2007; Gelineau-van Waes et al., 2008; Pickell et al., 2009; Salojin et al., 2011; Padmanabhan et al., 2013). Nevertheless, many of these mutations cause embryonic lethality by E10.5, a telltale sign of severe defects in placentation (Watson and Cross, 2005). The embryonic stage during which these phenotypes arise remains to be determined.

We previously reported the *Mtrr*^*gt*^ knock-down mutation generated through a gene-trap (gt) insertion into the mouse *Mtrr* locus (Elmore et al., 2007; Padmanabhan et al., 2013). The *Mtrr*^*gt*^ allele disrupts one-carbon metabolism by significantly reducing MTR activity (Elmore et al., 2007). Clinical indicators of human folate deficiency (Krishnaswamy and Madhavan Nair, 2001) are exhibited by *Mtrr*^*gt/gt*^ mice including plasma hyperhomocysteinaemia, macrocytic anaemia in adulthood, and a wide spectrum of developmental phenotypes at E10.5 including failure of neural tube closure (Elmore et al., 2007; Padmanabhan et al., 2013; Padmanabhan et al., 2018). Therefore, the *Mtrr*^*gt*^ mutation is a robust model to assess the effects of abnormal folate metabolism. Besides neural tube closure defects, other developmental phenotypes observed in *Mtrr*^*gt/gt*^ conceptuses at E10.5 are growth restriction, developmental delay, cardiac malformations, and placenta phenotypes whereby chorioallantoic attachment appear eccentric upon gross analysis or failed to occur (Padmanabhan et al., 2013). While the molecular cause and developmental timing of these phenotypes is currently unclear, alterations in genome-wide DNA methylation associated with gene misexpression were apparent in *Mtrr*^*gt/gt*^ conceptuses at E10.5 (Padmanabhan et al., 2013; Bertozzi et al., 2021; Blake et al., 2021), implicating an epigenetic mechanism.

Interestingly, the *Mtrr*^*gt*^ mouse line displays transgenerational epigenetic inheritance (TEI) of developmental phenotypes (Padmanabhan et al., 2013). TEI is a non-conventional mode of phenotype inheritance that is caused by an environmental stressor or metabolic disruption that is only present in the F0 generation (Blake and Watson, 2016). The mechanism of mammalian TEI is not well understood, though it is hypothesized that an epigenetic factor(s) is inherited via the germline to cause a phenotype in a manner that is independent of DNA base-sequence changes (Blake and Watson, 2016). An *Mtrr*^*gt*^ allele in a maternal grandparent (i.e., the F0 generation) was sufficient to disrupt germ cell DNA methylation (Blake et al., 2021) and cause growth defects and congenital malformations at E10.5 in the wildtype grandprogeny, in some cases up to the wildtype F4 generation (Padmanabhan et al., 2013). This phenomenon occurs when the parents (i.e., the F1 generation) and all subsequent generations are wildtype for *Mtrr* as determined through robust genetic pedigree analyses. Additionally, blastocyst transfer experiments determined that congenital malformations in the wildtype F2 grandprogeny were independent of the uterine environment of their wildtype F1 mothers (Padmanabhan et al, 2013). Thus, this evidence supports the occurrence of TEI in the *Mtrr*^*gt*^ model, though the specific mechanism is yet-to-be determined.

Here, we show evidence of developmental phenotypes in *Mtrr*^*gt/gt*^ conceptuses from E8.5, earlier than previously reported. We also observe a high prevalence of conceptuses with misaligned orientation within decidual swellings from E6.5. Misalignment occurred alongside twinning, which implicates folate metabolism in blastocyst orientation and spacing at implantation. Skewed orientation occurs along a spectrum with eccentric chorioallantoic attachment (ECA) now re-characterised as a severe form of misalignment. Furthermore, conceptus misalignment associates with developmental delay and heart defects, indicating that developmental progression of the embryo might be altered in this situation. Highly controlled genetic pedigree analysis and blastocyst transfer experiments reveal that severe conceptus misalignment is inherited transgenerationally in association with the *Mtrr* genotype of either maternal grandparent, and occurs independently of a uterine effect. Overall, we propose that defective folate metabolism disrupts blastocyst orientation and spacing over multiple generations, with implications for subsequent feto-placental development.

## MATERIALS AND METHODS

### Ethics

All experiments were performed in accordance with UK Home Office regulations under the Animals (Scientific Procedures) Act 1986 Amendment Regulations 2012 and underwent review by the University of Cambridge Animal Welfare and Ethical Review Body.

### Mice

The *Mtrr*^*gt*^ mouse line was generated as previously described (Padmanabhan et al., 2013). Briefly, a gene-trap (gt) vector was inserted into intron 9 of the *Mtrr* gene. Upon germline transmission, the *Mtrr*^*gt*^ allele was backcrossed to C57Bl/6J mice for eight generations (Padmanabhan et al., 2013). C57Bl/6J mice (The Jackson Laboratory) were used as controls and were bred in house separately from the *Mtrr*^*gt*^ mouse line. All mice were housed in a temperature- and humidity-controlled environment with a 12-hour light/dark cycle, and fed a standard chow (Rodent No. 3 chow, Special Diet Services, Essex, UK) *ad libitum* from weaning (includes folic acid at 2.99 mg/kg of diet, methionine at 0.37%, choline at 1422.4 mg/kg of diet, and vitamin B_12_ at 19.2 ug/kg of diet). *Mtrr*^*gt/gt*^ conceptuses at were derived from *Mtrr*^*gt/gt*^ intercrosses unless otherwise stated. For assessment of maternal effect, *Mtrr*^*+/+*^, *Mtrr*^*+/gt*^ and *Mtrr*^*gt/gt*^ female mice derived from *Mtrr*^*+/gt*^ intercrosses were mated with C57Bl/6J males, and the resulting litters were dissected at E6.5. For assessment of maternal grandparental effect, F0 *Mtrr*^*+/gt*^ male or female mice were crossed with a C57Bl/6J mate. Their F1 *Mtrr*^*+/+*^ daughters were mated with C57Bl/6J males to generate F2 *Mtrr*^*+/+*^ progeny for dissection at E10.5 or for littering out. F2 *Mtrr*^*+/+*^ females were mated with C57Bl/6J males to generate F3 *Mtrr*^*+/+*^ progeny for dissection at E10.5 or littering out. F3 *Mtrr*^*+/+*^ females were mated with C57Bl/6J males to generate F4 *Mtrr*^*+/+*^ conceptuses for dissection at E10.5.

### Genotyping

DNA samples were obtained from ear tissue or yolk sac for PCR genotyping of the *Mtrr*^*gt*^ allele or sex as reported previously (Padmanabhan et al., 2013; Tunster, 2017).

### Dissections and phenotyping

Noon of the day that the vaginal plug was detected was considered E0.5. Pregnant female mice were euthanized via cervical dislocation. Dissections were performed in cold 1x phosphate buffered saline (PBS). Each conceptus was individually scored for phenotypes (see below). Whole implantation sites at E6.5, or placentas and embryos separately at E8.5, E10.5, E14.5, and E18.5 were weighed, photographed, and either snap frozen in liquid nitrogen for molecular analysis or fixed in 10% natural-buffered formalin (Cat. No. HT501128, Sigma-Aldrich) at 4°C overnight and paraffin embedded for sectioning using standard protocols.

A rigorous phenotyping regime was performed as previously described (Padmanabhan et al., 2013). Briefly, conceptuses were defined as ‘phenotypically normal’ if they displayed 6-12 somite pairs at E8.5 and 30-40 somite pairs at E10.5 (according to e-Mouse Atlas Project; http://www.emouseatlas.org) and displayed no growth defect or gross abnormalities. For analysis at E10.5 only, an embryo was growth restricted if its crown-rump length was less than two standard deviations from the control C57Bl/6J mean crown-rump length. Growth restricted embryos also displayed normal somite pair counts and no other phenotype. At E8.5 and E10.5, developmentally delayed conceptuses exhibited fewer than six somite pairs or 30 somite pairs, respectively, but were otherwise normal for the developmental stage indicated by the somite counts. Severely affected conceptuses displayed one or more abnormality with appreciation for the developmental stage (e.g., somite pair count), as some of these embryos were developmentally delayed. Abnormalities included failure of neural tube closure and/or failure to initiate of embryo turning, heart looping defect, pericardial oedema, a placenta defect including eccentric or absent chorioallantoic attachment, or ‘twinning’ whereby two or more conceptuses shared a single implantation site. An implantation site that consisted of a decidual swelling containing an ill-defined mass of fetal-derived cells was categorised as a resorption and considered dead. Conceptuses were allocated a phenotypic category based on their most severe phenotype and were only counted once.

### Blastocyst transfer

Blastocyst transfer experiments were performed and analysed as previously described in detail (Padmanabhan et al., 2013). Briefly, using M2 media (Cat. No. M7167, Sigma-Aldrich), pre-implantation wildtype embryos were flushed at E3.25 from the uteri of *Mtrr*^*+/+*^ females (derived from one *Mtrr*^*+/gt*^ parent) mated with C57B/6J males. Females were not superovulated. After a brief embryo culture period to prepare the recipient females, embryos were surgically transferred into the uteri of E2.5 pseudopregnant B6D2F1 recipient females. C57Bl/6J embryos were similarly transferred as controls. Embryos were transferred regardless of appearance and litters were never pooled. Transferred embryos were dissected at E10.5, the timing of which corresponded to the staging of the recipient female (Ueda et al., 2003), and scored for phenotypes as described above.

### Immunohistochemistry

Whole implantation sites or placentas were carefully oriented in paraffin for transverse sectioning at 5 µm for conceptuses at E6.5 and 7 µm for conceptuses at E8.5. Tissue sections that were centrally located were selected for analysis, and deparaffinized in xylene (Cat. No. IS8126, Thermo Fisher Scientific) before rehydration in descending concentrations of ethanol. Hematoxylin (Cat No. MHS32, Sigma-Aldrich) and eosin (Cat No. 861006, Sigma-Aldrich) staining of histological sections was performed using standard protocols. For immunohistochemistry staining, endogenous peroxidase activity was quenched using 3% H_2_O_2_ (Cat No. H/1750/15, Thermo Fisher Scientific). Antigen retrieval was performed by treatment of tissue sections with porcine trypsin tablets (Cat No. T7168-20TAB, Sigma-Aldrich) for 10 min at room temperature. Tissue sections were first washed with 1x PBST (1x PBS with 20% Tween20) and then in blocking serum (5% serum (Cat. No. D9663, Sigma-Aldrich), 1% bovine serum albumin (BSA; Cat. No. A4503, Sigma-Aldrich) in 1x PBST) for 1 hour. Sections were exposed to primary antibodies diluted in blocking serum overnight at 4°C. The primary antibodies used and dilutions were: 1:300 rabbit polyclonal anti-COX2 (Cat. No. ab15191, abcam, Cambridge, UK, RRID:AB_2085144), 1:100 rabbit anti-MTRR (Cat. No. 26944-1-AP, Proteintech Europe, Manchester, UK, RRID:AB_2880694), and 1:150 rabbit anti-PGR (Cat. No. ab63605, abcam, RRID:AB 1142326). After thorough washing in 1x PBS-T, tissue sections were incubated for 1 hour at room temperature with donkey anti-rabbit IgG polyclonal antibody conjugated with horseradish peroxidase (Cat. No. ab6802, abcam, RRID:AB_955445) diluted to 1:300 in blocking serum. Peroxidase substrate reactions were conducted with 3,3’-diaminobenzidine (DAB) chromagen substrate kit according to the manufacturer’s instructions (Cat. No. ab64238, abcam). Tissue sections were counterstained with haematoxylin and coverslip mounted using DPX mountant (Cat. No. 360294, VWR).

### In situ hybridisation

In situ hybridisation probes were generated with gene-specific primer sets that contained T7 RNA polymerase promoter sequence (TAATACGACTCACTATAGGG) attached to each reverse primer and T3 RNA polymerase promoter sequence (AATTAACCCTCACTAAAGGG) attached to each forward primer (Outhwaite et al., 2019). See Supplementary Table 1 for gene specific promoter sequences. Probe synthesis of digoxigenin (DIG)-labelled anti-sense and sense probes was performed using DIG RNA labelling Mix (Cat. No. 11277073910, Roche, Welwyn Garden City, UK) according to the manufacturer’s instructions. The *in situ* hybridization protocol was performed as previously described in detail (Simmons et al., 2008) with the following alteration: paraffin-embedded tissue sections were used and therefore, a dewaxing and rehydration step was included.

### Quantitative reverse transcription PCR (RT-qPCR)

RNA was extracted from whole implantation sites at E6.5 using Trizol (Cat. No. T9424, Sigma-Aldrich) according to the manufacturer’s instructions. cDNA was synthesised using RevertAid H Minus reverse transcriptase (Cat. No. EP0452, Thermo Fisher Scientific) and random hexamer primer (Cat. No. SO142, Thermo Fisher Scientific) using 1-2 µg of RNA in a 20-µl reaction according to manufacturer’s instructions. PCR amplification was performed using MESA Green qPCR MasterMix for SYBR assay (Cat. No. RT-SY2X-03+WOUN, Eurogentec, Ltd.) on a DNA Engine Opticon2 thermocycler (BioRad). Experiments were conducted with technical duplicates or triplicates and using at least four biological replicates. Transcript levels were normalised to *Polr2a* (Solano et al., 2016) and analysed using the ΔΔCt method (Livak and Schmittgen, 2001). Transcript levels in C57Bl/6J tissue were normalized to 1. For primer sequences, refer to Supplementary Table 1.

### Statistics

Statistical analyses were performed using GraphPad Prism 8 software (La Jolla, CA, USA). Parametric data was analysed by unpaired t test or one-way ANOVA. Non-parametric data was analysed by Mann Whitney test. Correlations were analysed by linear regression or Fisher’s exact test. P<0.05 was considered significant.

### Software and imaging

Primer sequences were designed using the NCBI primer design tool (Ye et al., 2012). Conceptuses were dissected using a Zeiss SteReo Discovery V8 microscope and photographed with an AxioCam MRc5 camera and AxioVision 4.7.2 software (Carl Zeiss). Tissue sections were scanned using NanoZoomer (Hamamatsu Photonics, UK) and viewed using NDP Scan 2.7 software (Hamamatsu Photonics). Images were analysed using ImageJ software (NIH, Bethesda, MD, USA). Graphs were generated using Graphpad Prism 8 software.

## RESULTS

### *Mtrr*^*gt/gt*^ conceptuses exhibit developmental phenotypes at E8.5

MTRR protein was widely expressed in wildtype embryonic and extra-embryonic cell types including the yolk sac, amnion, allantois, and trophoblast progenitors (i.e., the chorion and EPC) at E8.5 (Figures 1A-C). This expression pattern was consistent with other folate metabolic enzymes and transporters at E8.5 (Cherukad et al., 2012). While expressed throughout the mesometrial and lateral decidua (Figure 1A), the antimesometrial decidua displayed striped MTRR expression in the primary and secondary decidual zones at E8.5 (Figure 1C) suggesting regionalised expression in the decidua. Overall, the broad expression of MTRR protein in wildtype conceptuses at E8.5 suggested a potential mechanistic role of folate metabolism in early fetoplacental development and decidualisation.

**Figure 1.**
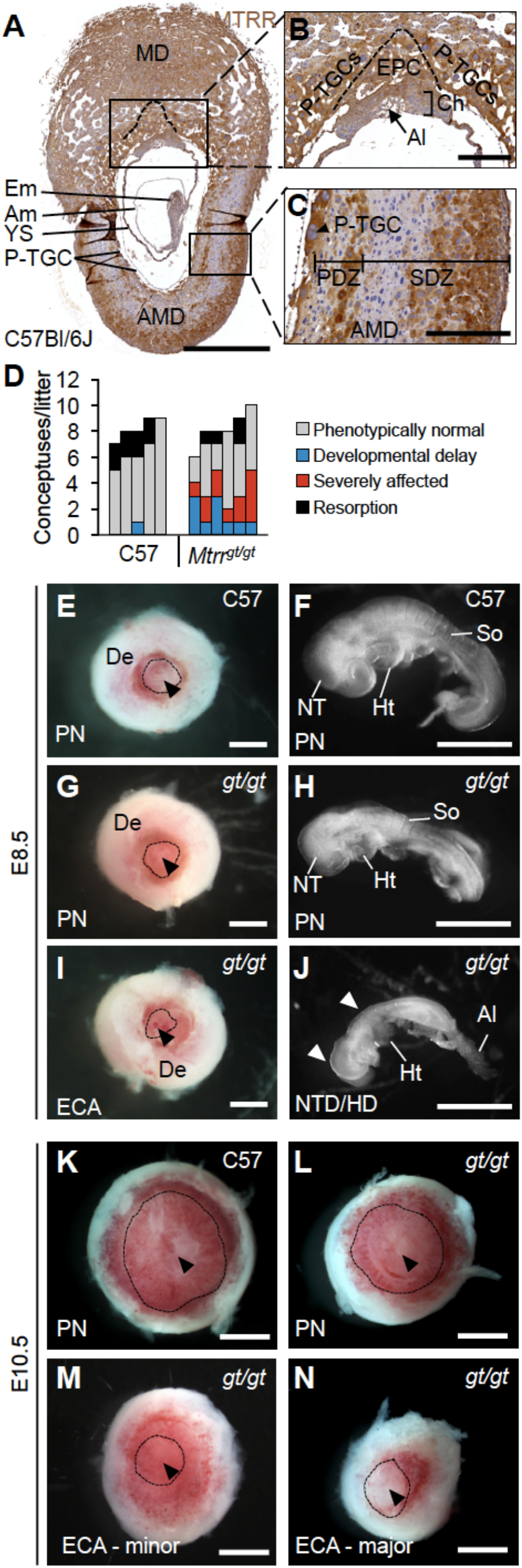
Wide spectrum of phenotypes was observed in *Mtrr*^*gt/gt*^ mouse conceptuses at E8.5. (A-C) MTRR protein expression (brown) in a control wildtype C57Bl/6J implantation site at E8.5. DNA, blue. Boxed regions in (A) shown at higher magnification in (B) and (C). N=5 implantation sites, 3 technical replicates. Scale bars: (A) 1 mm, (B-C) 250 µm. (D) Graph depicting the frequency of phenotypes per litter in conceptuses at E8.5 from C57Bl/6J control crosses and *Mtrr*^*gt/gt*^ intercrosses. Each bar represents one litter. Grey, phenotypically normal; black, resorption; blue, developmental delay (<6 somite pairs); red, severely affected. See also Table 1. (E-J) Images of conceptuses captured at E8.5 depicting (E-F) phenotypically normal (PN) C57Bl/6J (C57) control placenta and embryo, and (G-H) phenotypically normal and (I-J) severely affected *Mtrr*^*gt/gt*^ (*gt/gt*) conceptuses. The *Mtrr*^*gt/gt*^ placenta in (I) displays eccentric chorioallantoic attachment (ECA). The *Mtrr*^*gt/gt*^ embryo in (J) displays normal developmental stage (6 somite pairs) yet initiation of neural tube closure defects (NTD) and heart defect (HD). White arrowheads indicate open regions of the neural tube. The pairwise embryos and placentas at E8.5 depicted are not from the same conceptuses. (K-N) Images of placentas captured at E10.5 from (K) a C57Bl/6J control conceptus, (L) a phenotypically normal *Mtrr*^*gt/gt*^ conceptus, (M-N) *Mtrr*^*gt/gt*^ conceptuses exhibiting eccentric chorioallantoic attachment with (M) minor or (N) major severity. (E, G, I, K-N) Dotted line, outline of chorion; arrowhead, region of allantois attachment. Scale bars, E, G, I, K-L: 1 mm; F, H, J: 0.5 mm. Al, allantois; Am, amnion; AMD, antimesometrial decidua; Ch, chorion; De, decidua; ECA, eccentric chorioallantoic attachment; Em, embryo proper; EPC, ectoplacental cone (dotted line in A-B); He, head; Ht, heart; MD, mesometrial decidua; NT, neural tube; PDZ, primary decidual zone; PN, phenotypically normal; P-TGCs, parietal trophoblast giant cells; SDZ, secondary decidual zone; So, somite; YS, yolk sac.

To explore whether phenotypes previously observed in the *Mtrr*^*gt*^ mouse line at E10.5 were present at an earlier time point, *Mtrr*^*gt/gt*^ conceptuses derived from *Mtrr*^*gt/gt*^ intercrosses were rigorously scored for gross abnormalities at E8.5 (N=50 conceptuses from 6 litters). C57Bl/6J conceptuses, which are genetically wildtype for *Mtrr* and bred separately from the *Mtrr*^*gt*^ mouse line, were used as controls (N=43 conceptuses from 5 litters) because the *Mtrr*^*gt*^ allele was backcrossed into the C57Bl/6J mouse strain and because *Mtrr*^*+/+*^ conceptuses derived from *Mtrr*^*+/gt*^ intercrosses display multigenerational inheritance of developmental phenotypes (Padmanabhan et al., 2013). Similar to E10.5 (Padmanabhan et al., 2013), *Mtrr*^*gt/gt*^ litter sizes and resorption rates were consistent with controls at E8.5 (Figure 1D; Table 1). Allocation to a phenotypic category depended upon the most severe phenotype exhibited by each conceptus (see methods). The majority of C57Bl/6J conceptuses at E8.5 were phenotypically normal (79.1% of conceptuses), displaying 6-12 somite pairs and no other abnormalities (Figures 1D-F; Table 1). Nearly all other C57Bl/6J conceptuses were resorbed (18.6% of conceptuses; Figure 1D; Table 1), appearing as decidual swellings with an ill-defined mass of embryo-derived cells. In contrast, only 48.0% of *Mtrr*^*gt/gt*^ conceptuses at E8.5 were phenotypically normal upon gross inspection (Figures 1D,G,H; Table 1). The remaining *Mtrr*^*gt/gt*^ conceptuses were either resorbed (8.0% of conceptuses; Figure 1D; Table 1), or exhibited one or more developmental phenotype (Figures 1D,I,J; Table 1). These phenotypes included developmental delay, defined as embryos with <6 somite pairs yet otherwise normal for the developmental stage (20.0% of conceptuses; p<0.05; Figure 1D; Table 1), or severe abnormalities (24.0% of conceptuses; p<0.01; Figures 1D,I,J; Table 1). Defects that were considered severe were failure of neural tube closure (Figure 1J), a lack of initiation of turning, abnormal heart looping despite normal somite pair count, pericardial edema (Figure 1J), and/or eccentric (Figure 1I) or absent chorioallantoic attachment. The most frequent severe abnormalities in *Mtrr*^*gt/gt*^ conceptuses at E8.5 were chorioallantoic attachment defects (16.0% of conceptuses; Table 1) and failure of neural tube closure (14.0% of conceptuses; Table 1). In contrast, C57Bl/6J conceptuses at E8.5 infrequently displayed developmental delay (2.3% of conceptuses; Figure 1D; Table 1) and severe abnormalities were undetected (Figure 1D; Table 1). These data were similar to *Mtrr*^*gt/gt*^ conceptuses at E10.5 (Padmanabhan et al., 2013), and suggested that the developmental phenotypes caused by the *Mtrr*^*gt*^ mutation likely originate earlier than E8.5.

**Table 1.** Frequency of developmental phenotypes in *Mtrr*^*gt/gt*^ conceptuses at E8.5

| Embryonic genotype | C57Bl/6 | *Mtrr<sup>gt/gt</sup>* |
| --- | --- | --- |
| **No. of conceptuses**<br>(No. of litters) | **43**<br>(5) | **50**<br>(6) |
| **No. of conceptuses/litter**<br>[mean ± sd] | **8.6 ± 0.2** | **8.3 ± 0.6** |
| **Phenotypically Normal**<br>[6-12 somite pairs] | **6.8 ± 0.7**<br>(79.1%) | **4.0 ± 0.7\***<br>(48.8%) |
| **Developmentally Delay**<br>[<6 somite pairs] | **0.2 ± 0.2**<br>(2.3%) | **1.7 ± 0.4\***<br>(20.0%) |
| **Severe abnormalities** | **0.0 ± 0.0**<br>(0.0 %) | **2.0 ± 0.4\*\***<br>(24.0%) |
| >1 severe abnormality | **0.0 ± 0.0**<br>(0.0%) | **0.05 ± 0.06**<br>(6.0%) |
| **Placental defects** |  |  |
| Off-centre chorioallantoic attachment | **0.0 ± 0.0**<br>(0.0%) | **0.11 ± 0.13**<br>(12.0%) |
| No chorioallantoic attachment | **0.0 ± 0.0**<br>(0.0%) | **0.05 ± 0.07**<br>(4.0%) |
| **Embryonic defects** |  |  |
| Heart not looped | **0.0 ± 0.0**<br>(0.0%) | **0.02 ± 0.05**<br>(2.0%) |
| Neural tube defects | **0.0 ± 0.0**<br>(0.0%) | **0.13 ± 0.11\***<br>(14.0%) |
| **Resorption** | **1.6 ± 0.4**<br>(18.6%) | **0.7 ± 0.3**<br>(8.0%) |

Data displayed as mean incidence per litter ± sd. Percentage of all conceptuses assessed in brackets. \*P<0.05; \*\*P<0.01

### A low frequency of twinning occurs in *Mtrr*^*gt/gt*^ conceptuses throughout gestation

Twinning in mice occurs when at least two conceptuses occupy a single decidual swelling. We observed a low frequency of dizygotic twinning in the *Mtrr*^*gt*^ mouse line at several developmental stages (0.7-2.3% of implantation sites, N=118-308 sites depending on the stage assessed) (Figures 2A,B). Twinning was never detected in C57Bl/6J controls (N=314 conceptuses). The occurrence of male and female embryos together in a single implantation site indicated dizygosity and suggested that twinning in the *Mtrr*^*gt*^ mouse line resulted from poor blastocyst spacing at implantation. Since *Mtrr*^*gt/gt*^ implantation sites were otherwise fairly evenly spaced along the length of *Mtrr*^*gt/gt*^ uteri (Figure 2C) and no more than one twinned site was present per *Mtrr*^*gt/gt*^ litter, a uterine-specific defect was unlikely. Therefore, we hypothesise that abnormal folate metabolism disrupts spacing of some blastocysts at implantation via an embryo-specific defect.

**Figure 2.**
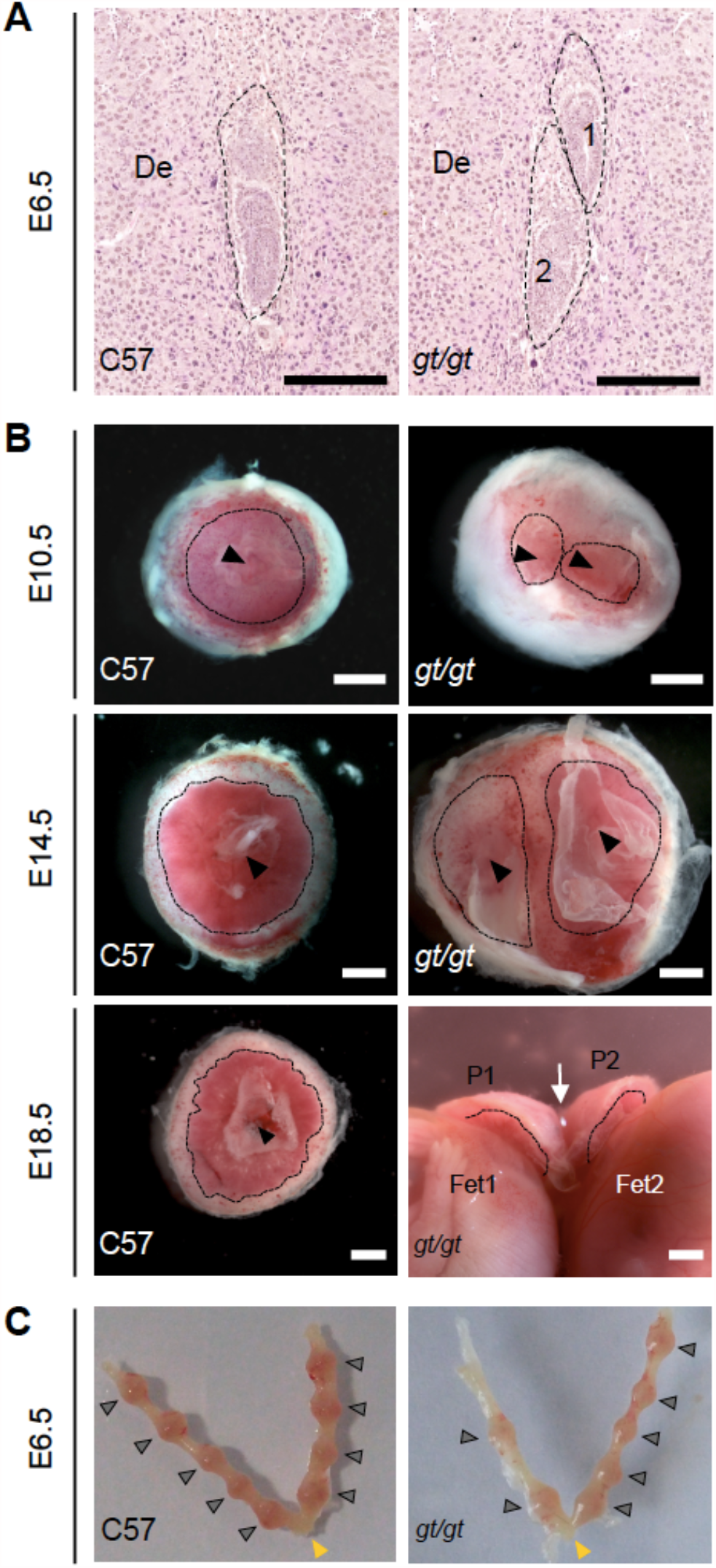
Twinning is observed at a low frequency in *Mtrr*^*gt/gt*^ mouse conceptuses throughout development. (A-B) Images depicting twinning in *Mtrr*^*gt/gt*^ (*gt/gt*) conceptuses at several stages of development. (A) Histological sections of a C57Bl/6J (C57) implantation site at E6.5 with a single conceptus (dashed outline, left-hand image) and of an *Mtrr*^*gt/gt*^ implantation site at E6.5 with two conceptuses (dashed outlines, right-hand image). Scale bar, 250 µm. (B) Whole placentas from single C57Bl/6J (C57) conceptuses (left-hand images) and twinned *Mtrr*^*gt/gt*^ (*gt/gt*) conceptuses (right-hand images) at E10.5, E14.5 and E18.5 as viewed on the chorionic plate. Black dotted line, base of labyrinth; black arrowhead, allantois attachment site; white arrow indicates where the placentas at E18.5 are joined. De, decidua; P, placenta; Fet, fetus. Scale bars, 1 mm. (C) Dissected uteri at E6.5 from C57Bl/6J (C57) and *Mtrr*^*gt/gt*^ (*gt/gt*) females. Grey arrowheads, individual implantation sites. Yellow arrowhead, cervix.

### Eccentric chorioallantoic attachment is caused by skewed orientation of the entire conceptus

Our initial focus was placed on the chorioallantoic attachment defects, due to the high frequency in the *Mtrr*^*gt*^ model. While no chorioallantoic attachment defects were detected in C57Bl/6J conceptuses at E8.5 or E10.5 (Table 1, Figures 1E,K, 6A) (Padmanabhan et al., 2013), *Mtrr*^*gt/gt*^ conceptuses showed two distinct chorioallantoic attachment phenotypes: attachment failure (E8.5: 4.0% of conceptuses; E10.5: 0.3% of conceptuses) and eccentric chorioallantoic attachment (ECA) (E8.5: 12.0% of conceptuses; E10.5: 5.8% of conceptuses; Table 1, Figures 1I,L-N, 6B) (Padmanabhan et al., 2013). Declining phenotypic frequencies during gestation suggested that some *Mtrr*^*gt/gt*^ conceptuses with a chorioallantoic attachment defect likely undergo lethality between E8.5 and E10.5.

To better understand the ECA phenotype, centrally located transverse histological sections of C57Bl/6J and *Mtrr*^*gt/gt*^ placentas were examined at E8.5 after completion of chorioallantoic attachment (Cross et al., 2003). We expected to observe allantoic attachment at the side of the chorion rather than the centre. However, among the *Mtrr*^*gt/gt*^ placentas, a proportion of the trophoblast compartments were noticeably askew within their decidual swellings (Figures 3A-C). The degree of skewed growth was determined by measuring the angle between the midline of the decidual swelling and the midline of the chorion or EPC (Figures 3A-C). In *Mtrr*^*gt/gt*^ placentas, the angles of chorion and EPC skewing were proportionate with an average angle of 10.2° ± 10.5° (mean ± s.d.) and 8.3° ± 15.9°, respectively (Figures 3D-F). These values showed an increased trend compared to C57Bl/6J conceptuses (chorion: 2.4° ± 4.5° (Mann-Whitney test, P=0.020); EPC: 0.3° ± 1.0°, P=0.143; Figures 3D-E). Given the proportionality of chorion and EPC misalignment (Figure 3F), we hypothesised that the skewed orientation extended to the entire *Mtrr*^*gt/gt*^ conceptus. Inter-individual variability of chorion/EPC skewing between *Mtrr*^*gt/gt*^ conceptuses was apparent (Figures 3D,E) suggesting skewing occurred along a spectrum of severity.

**Figure 3.**
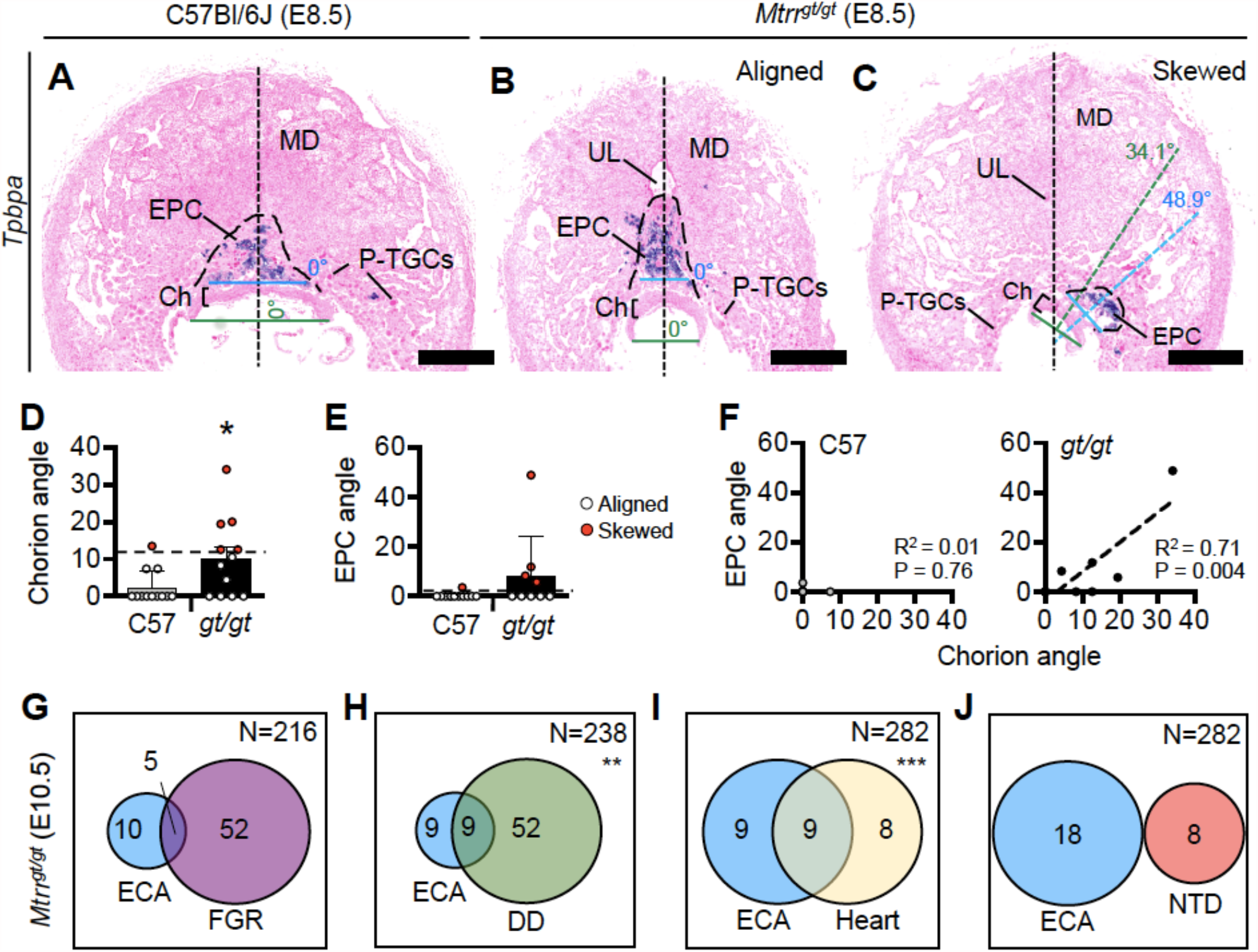
*Mtrr*^*gt/gt*^ mouse conceptuses at E8.5 demonstrate misaligned orientation with incomplete penetrance. (A-C) Histological sections of placentas at E8.5 from (A) a C57Bl/6J conceptus and (B-C) *Mtrr*^*gt/gt*^ conceptuses with (B) aligned or (C) skewed orientation. Ectoplacental cone (EPC, dashed line) is depicted by *Tpbpa* mRNA expression (purple) as determined via *in situ* hybridisation. The lines bisecting the chorion (dotted green line) or EPC (dotted blue line) are shown. The angle between each of these lines and the line bisecting the placenta (dotted black line) was determined for each placenta. Scale bar = 500 µm. Ch, chorion; EPC, ectoplacental cone; MD, mesometrial decidua; P-TGCs, parietal trophoblast giant cells; UL, uterine lumen remnant. (D-E) Data showing the average (mean ± sd) (D) chorion angle or (E) EPC angle in C57Bl/6J (C57) controls and *Mtrr*^*gt/gt*^ (*gt/gt*) conceptuses at E8.5 (N=9-12 conceptuses/group). Conceptuses with chorion or EPC angles >2 sd above the control mean (black dashed line [chorion = 11.4°; EPC = 2.4°]) were defined as skewed (red dots). Two-tailed Mann-Whitney test, *P=0.028. (F) Linear regression analysis between chorion and EPC angles for C57Bl/6 conceptuses (C57; N=10 conceptuses) and *Mtrr*^*gt/gt*^ conceptuses (*gt/gt*; N=9 conceptuses). (G-J) Venn diagrams depicting overlap between eccentric chorioallantoic attachment (ECA; severe skewing) in *Mtrr*^*gt/gt*^ conceptuses at E10.5 and other phenotypes including (G) fetal growth restriction (FGR), (H) developmental delay (DD), (I) heart malformations, and (J) neural tube defects (NTDs) (see methods). N values in top right indicate total number of *Mtrr*^*gt/gt*^ conceptuses assessed. Fisher’s exact test, **P<0.01, ***P<0.001.

Therefore, we defined a chorion or EPC as ‘skewed’ when its angle was >2 s.d. above the control mean (chorion: >11.4°; EPC: >2.4°; Figures 3D,E). Using this definition, 50.0% (6/12) of *Mtrr*^*gt/gt*^ trophoblast compartments at E8.5 were skewed compared to only 14.3% (2/14) in C57Bl/6J conceptuses (Figures 3D,E). We propose that the ECA phenotype identified during gross dissection likely denotes the extreme end of the skewed conceptus orientation spectrum. Moreover, less severe conceptus skewing was likely unappreciated during gross dissection.

### Embryonic phenotypes associate with skewed orientation in *Mtrr*^*gt/gt*^ conceptuses at E10.5

To determine whether embryonic phenotypes were associated with conceptus misalignment, we assessed our phenotyping data from *Mtrr*^*gt/gt*^ conceptus dissections at E10.5 (Padmanabhan et al., 2013; Padmanabhan et al., 2017) for overlap between ECA (representing severe skewing) and embryo growth defects or congenital malformations. We selected E10.5 for assessment due to the size of this data set (>215 *Mtrr*^*gt/gt*^ conceptuses at E10.5) and because the detrimental effects of misaligned orientation might be more apparent at this stage. A caveat of this analysis was that only severe conceptus skewing was assessed using these criteria. While ECA did not associate with fetal growth restriction (5/15 *Mtrr*^*gt/gt*^ embryos with 30-40 somite pairs had ECA and crown-rump lengths <2 s.d. of control mean, Fisher’s exact test, P=0.193; Figure 3G), there was a correlation with developmental delay (9/18 *Mtrr*^*gt/gt*^ embryos with <30 somite pairs had ECA, Fisher’s exact test, P=0.003; Figure 3H). Separately, we observed that ECA associated with heart malformations including pericardial oedema or reversed heart looping (9/18 of *Mtrr*^*gt/gt*^ embryos had ECA and a heart abnormality; Fisher’s exact test, P<0.0001; Figure 3I). Yet, zero out of 18 *Mtrr*^*gt/gt*^ embryos that exhibited ECA failed to close their neural tube (Figure 3J). While a causal relationship is yet-to-be established, it is possible that skewed conceptus orientation influenced subsequent embryo development.

### Skewed orientation does not alter *Mtrr*^*gt/gt*^ chorion patterning at E8.5

To explore whether skewed conceptus orientation associated with abnormal development of the trophoblast population, in situ hybridization was performed to detect key chorion and EPC trophoblast marker genes (Simmons et al., 2008). Histological sections of C57Bl/6J control (N=6 placentas) and *Mtrr*^*gt/gt*^ placentas (N=14 placentas) were assessed at E8.5, with *Mtrr*^*gt/gt*^ placentas subdivided into those with aligned and skewed orientation. Trophoblast marker gene expression of *Tpbpa* (EPC marker), *Hand1* (sinusoidal TGC progenitor and P-TGCs), *Syna* (syncytiotrophoblast-I progenitors) and *Gcm1* (syncytiotrophoblast-II progenitors) was present in all *Mtrr*^*gt/gt*^ placentas assessed, regardless of the degree of skewing (Figures 3A-C, 4A-I). These data indicated that the trophoblast populations were present and patterned in *Mtrr*^*gt/gt*^ placentas, and thus not greatly affected by skewed conceptus growth. However, *Prl7b1* mRNA expression (invasive trophoblast cell marker) was substantially reduced only in *Mtrr*^*gt/gt*^ placentas with skewed orientation. The few *Prl7b1+* cells that were present were located in the lateral decidua rather than in the mesometrial decidua (Figures 4J-L). Mislocalisation of invasive trophoblast cells might have significant implications for uterine spiral artery remodelling and should be explored in future studies.

**Figure 4.**
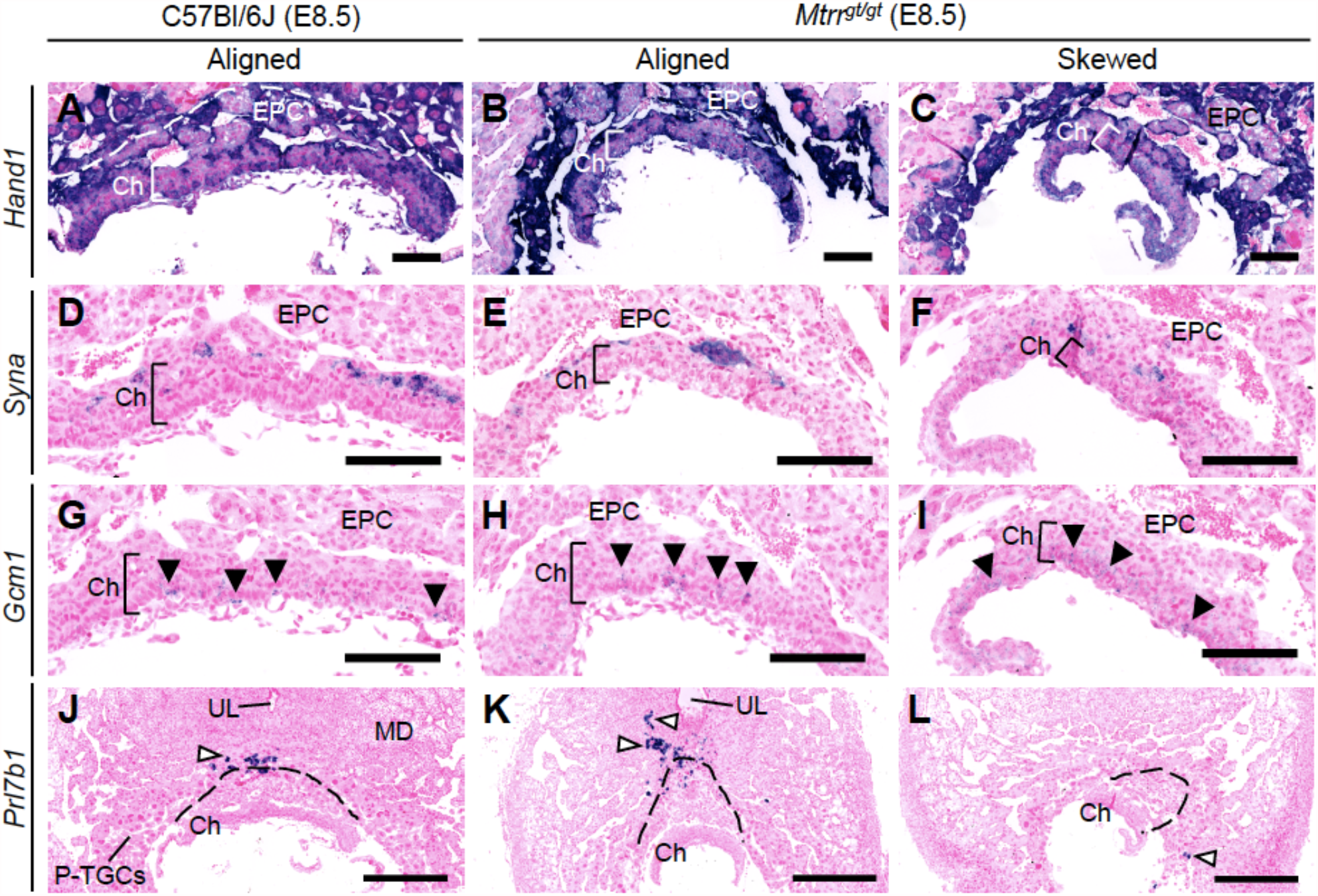
Patterning of the trophoblast lineages proceeds in *Mtrr*^*gt/gt*^ mouse conceptuses regardless of misalignment. Analysis of trophoblast gene marker expression (purple) via *in situ* hybridization in C57Bl/6J (N=6) and *Mtrr*^*gt/gt*^ placentas (N=14) at E8.5. *Mtrr*^*gt/gt*^ placentas with aligned and skewed orientation are shown. Trophoblast markers included (A-C) *Hand1* (widespread trophoblast marker including sinusoidal TGC progenitors of the apical chorion), (D-F) *Syna* (syncytiotrophoblast-I progenitors), (G-I) *Gcm1* (syncytiotrophoblast-II progenitors; branch point sites indicated by *Gcm1+* cells [black arrowheads]), and (J-L) *Prl7b1* (spiral artery TGCs, white arrowheads; dotted line indicates ectoplacental cone). Scale bars: (A-C) 1 mm, (D-L) 125 µm. Ch, chorion; EPC, ectoplacental cone; MD, mesometrial decidua; P-TGCs, parietal trophoblast giant cells; UL, uterine lumen remnant.

### *Mtrr*^*gt/gt*^ conceptus misalignment is not associated with decidual blood sinus area at E8.5

Another model of skewed EPC growth exhibits dilated decidual blood sinuses (Winterhager et al., 2013). This association suggested that defects in the maternal decidua might lead to skewed orientation of conceptus growth. Therefore, to investigate a similar link in *Mtrr*^*gt/gt*^ conceptuses at E8.5, combined blood sinus area in the lateral and mesometrial decidual regions was measured in centrally located transverse histological sections of C57Bl/6 and *Mtrr*^*gt/gt*^ implantation sites at E8.5 (N=6-12 placentas/group). Both aligned and skewed *Mtrr*^*gt/gt*^ conceptuses were separately considered. The average proportion of the decidua attributed to blood sinuses was similar in C57Bl/6J, *Mtrr*^*gt/gt*^ aligned, and *Mtrr*^*gt/gt*^ skewed placentas (P=0.259; Figures 5A-D). Furthermore, there was no correlation between decidual blood sinus area and chorion angle (P>0.30; Supplementary figure 2A) or EPC angle (P>0.14; Figure 5E) in either *Mtrr*^*gt/gt*^ group assessed compared to controls. Next, we examined whether left-right asymmetry in blood sinus area was evident and potentially guided the direction of conceptus misalignment. To do this, the ratio of right:left decidua blood sinus area was calculated. Similar to controls, decidual blood sinus area was similar on the right and left sides of *Mtrr*^*gt/gt*^ implantation sites, and was independent of whether conceptus skewing occurred (P=0.605; Figure 5F). For each implantation site, the angle and direction of conceptus skewing was then compared to right-left decidual blood sinus ratio. Using histological sections of individual implantation sites, the chorion or EPC angle was classified as positive if it was skewed to the right and negative if it was skewed to the left. This value was plotted against the ratio of right:left decidual blood sinus area for that site (Figure 5G, Supplementary figure 2B). In this model, a line-of-best-fit sloping downwards towards the right would indicate that the chorion/EPC angled away from the greater blood sinus area, and a slope upwards towards the right would indicate that the chorion/EPC angled towards the greater blood sinus area. In C57Bl/6J controls, chorion and EPC skewing was absent regardless of whether the blood sinus area was greater on the right or left (Figure 5G, Supplementary figure 2B). Similarly in *Mtrr*^*gt/gt*^ conceptuses, there was no statistically significant relationship between blood sinus size and the direction of chorion or EPC growth (P>0.37; Figure 5G, Supplementary figure 2B). These data suggested that the volume of maternal blood perfused into the implantation site was unlikely related to the orientation of *Mtrr*^*gt/gt*^ conceptuses at E8.5.

**Figure 5.**
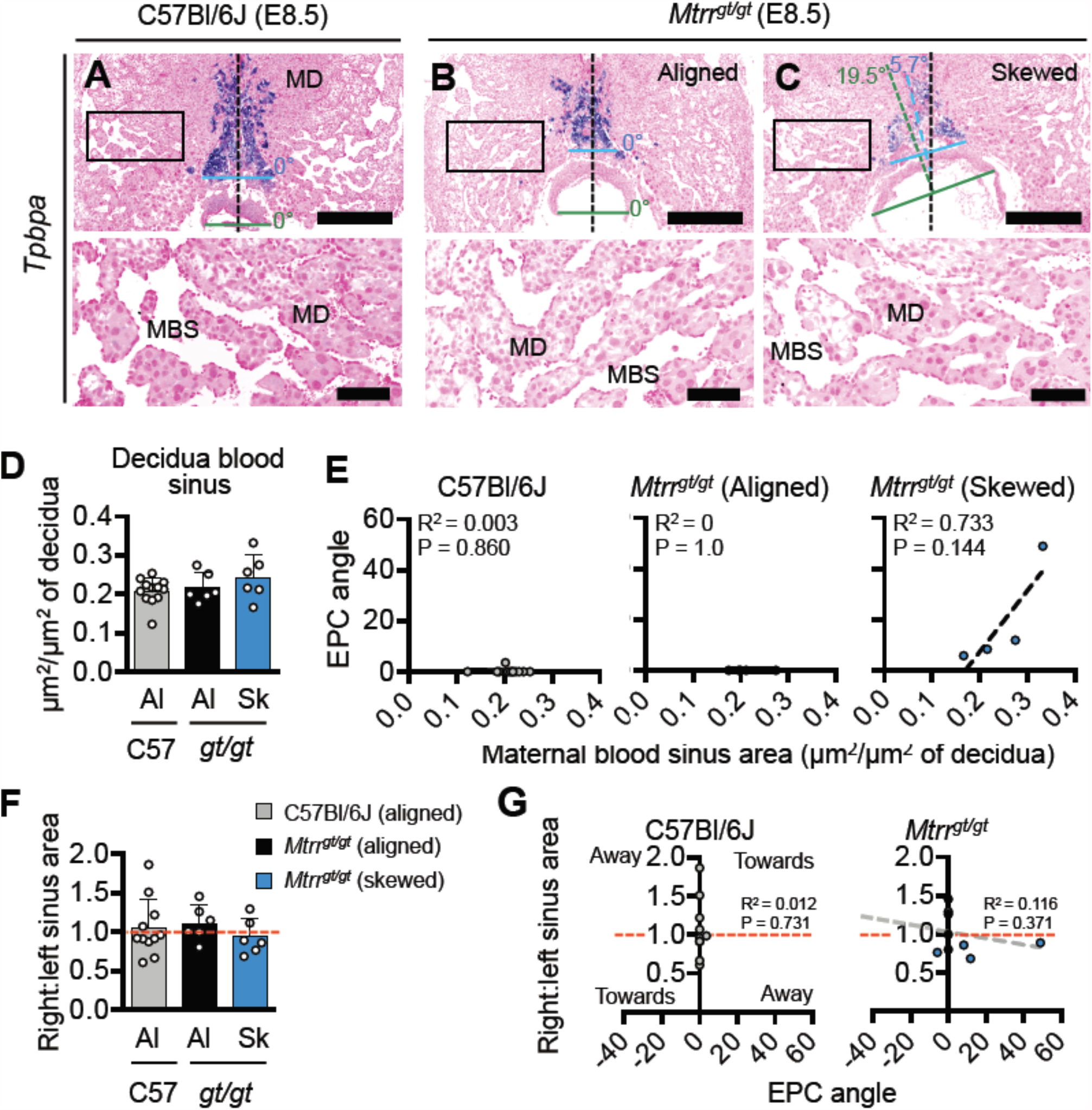
Mouse conceptus skewing does not correlate with decidual blood sinus area. (A-C) Histological sections of (A) C57Bl/6J and (B-C) *Mtrr*^*gt/gt*^ placentas at E8.5. Ectoplacental cones are stained for *Tpbpa* via *in situ* hybridization. DNA, pink. Boxed area indicates region of decidua shown at higher magnification in the image below. Placentas with (A-B) aligned and (C) skewed orientation are shown. N=6-12 placentas/group. Black dashed line bisects the decidua. Blue dashed line bisects the EPC. Green dashed line bisects the chorion. Skewing angles are indicated for chorion (green) and EPC (blue). MD, mesometrial decidua; MBS, maternal blood sinus in decidua. Scale bars: top panel, 500 µm; bottom panel, 100 µm. (D) Graph showing average area of decidual blood sinuses (mean ± sd) per total decidua area in histological sections of C57Bl/6J (grey bar) and *Mtrr*^*gt/gt*^ (black and blue bars) placentas at E8.5. Aligned (Al) and skewed (Sk) placentas are represented (N=6-12 placentas*/*group, with at least 3 sections assessed per placenta). One-way ANOVA, P=0.2586. (E) Linear regression analyses of decidua blood sinus area in relation to EPC angle for C57Bl/6J and *Mtrr*^*gt/gt*^ conceptuses. Aligned and skewed *Mtrr*^*gt/gt*^ placentas were considered separately (N=6-12 placentas*/*group). (F) Data depicting the ratio of decidua blood sinus area on the right and left sides of implantation sites from C57Bl/6J (C57) and *Mtrr*^*gt/gt*^ (*gt/gt*) conceptuses at E8.5. Data is represented as mean ± sd. Aligned (Al) and skewed (Sk) placentas were considered separately (N=6-12 placentas*/*group). Dashed red line indicates a ratio of 1. One-way ANOVA, P=0.6050. (G) To determine whether conceptus skewing occurred towards or away from the greatest decidual blood sinus area, linear regression analyses were performed between the EPC angle and the ratio of right:left decidua blood sinus area in C57Bl/6J and *Mtrr*^*gt/gt*^ conceptuses at E8.5 (N=6-12 conceptuses*/*group). Aligned (grey or black dots) and skewed (blue dots) conceptuses are shown. Dashed red line, right:left blood sinus area ratio of 1. Dashed grey line, line of best fit. See also Supplementary figures 2A-B.

### Conceptus skewing is associated with the *Mtrr* genotype of the maternal grandparents

To further consider whether skewed orientation in *Mtrr*^*gt/gt*^ conceptuses was caused by a maternal effect, we manipulated the maternal *Mtrr* genotype in highly controlled genetic pedigrees and assessed the frequency of conceptus skewing. *Mtrr*^*+/+*^, *Mtrr*^*+/gt*^, and *Mtrr*^*gt/gt*^ females were generated from *Mtrr*^*+/gt*^ intercrosses and mated to C57Bl/6J control males. Implantation sites were dissected at E6.5 (N=31-41 conceptuses/maternal genotype from 4 litters) and assessed for conceptus skewing and defects in decidualisation (see below). C57Bl/6J intercrosses were used as controls. The average litter size was significantly different between the pedigrees (Figure 6A; One-way ANOVA, P=0.0351) with greatest difference between litters from *Mtrr*^*+/gt*^ females (10.3 ± 1.0 conceptuses/litter [mean ± s.d.]) and *Mtrr*^*gt/gt*^ females (7.8 ± 1.7 conceptuses/litter, Dunn’s multiple comparison, P<0.05). To assess the extent of conceptus skewing caused by the maternal *Mtrr*^*gt*^ genotype at E6.5, centrally located transverse histological sections of implantation sites at E6.5 were examined (N=7-10 implantation sites/group). In a manner similar to the analysis at E8.5 (Figures 3A-C), the angle between the midlines of the implantation site and the entire conceptus was measured (Figures 6B,C). Most C57Bl/6J conceptuses were aligned along the mesometrial-antimesometrial axis with an average angle of 0.3° ± 0.9° (mean ± s.d.; Figures 6B,C). In contrast, conceptuses derived from *Mtrr*^*+/+*^, *Mtrr*^*+/gt*^ and *Mtrr*^*gt/gt*^ females showed average angles that were greater than controls (*Mtrr*^*+/+*^ mothers: 4.7° ± 5.8°; *Mtrr*^*+/gt*^ mothers: 4.2° ± 5.7°; *Mtrr*^*gt/gt*^ mothers: 6.2° ± 6.7°; Figure 6B). Statistical significance was not reached due to high inter-individual variability (Figure 6B). However, similar to our observations in *Mtrr*^*gt/gt*^ conceptuses at E8.5 (Figures 3D,E), skewing at E6.5 (regardless of maternal *Mtrr* genotype) occurred across a spectrum of severity (Figure 6B). Therefore, we similarly defined a conceptus at E6.5 as ‘skewed’ when it had an angle >2 s.d. above the control mean (i.e., >2.2°). Using this definition, only 11% (1/9) of C57Bl/6J conceptuses were skewed compared to 60% (6/10), 44% (4/9), and 57% (4/7) of conceptuses with *Mtrr*^*+/+*^, *Mtrr*^*+/gt*^, and *Mtrr*^*gt/gt*^ mothers, respectively (Figure 6B). Altogether, these data provided evidence that skewed conceptus growth was established earlier than E6.5, potentially at implantation. Furthermore, there was a similar frequency of misaligned conceptuses derived from *Mtrr*^*+/+*^ mothers as from *Mtrr*^*+/gt*^ and *Mtrr*^*gt/gt*^ mothers, implicating the *Mtrr*^*+/gt*^ maternal grandparents in the inheritance of this phenotype. Multigenerational inheritance of developmental phenotypes was previously reported in the *Mtrr*^*gt*^ model in other contexts (Padmanabhan et al., 2013; Padmanabhan et al., 2017; Padmanabhan et al., 2018) and will be explored below.

**Figure 6.**
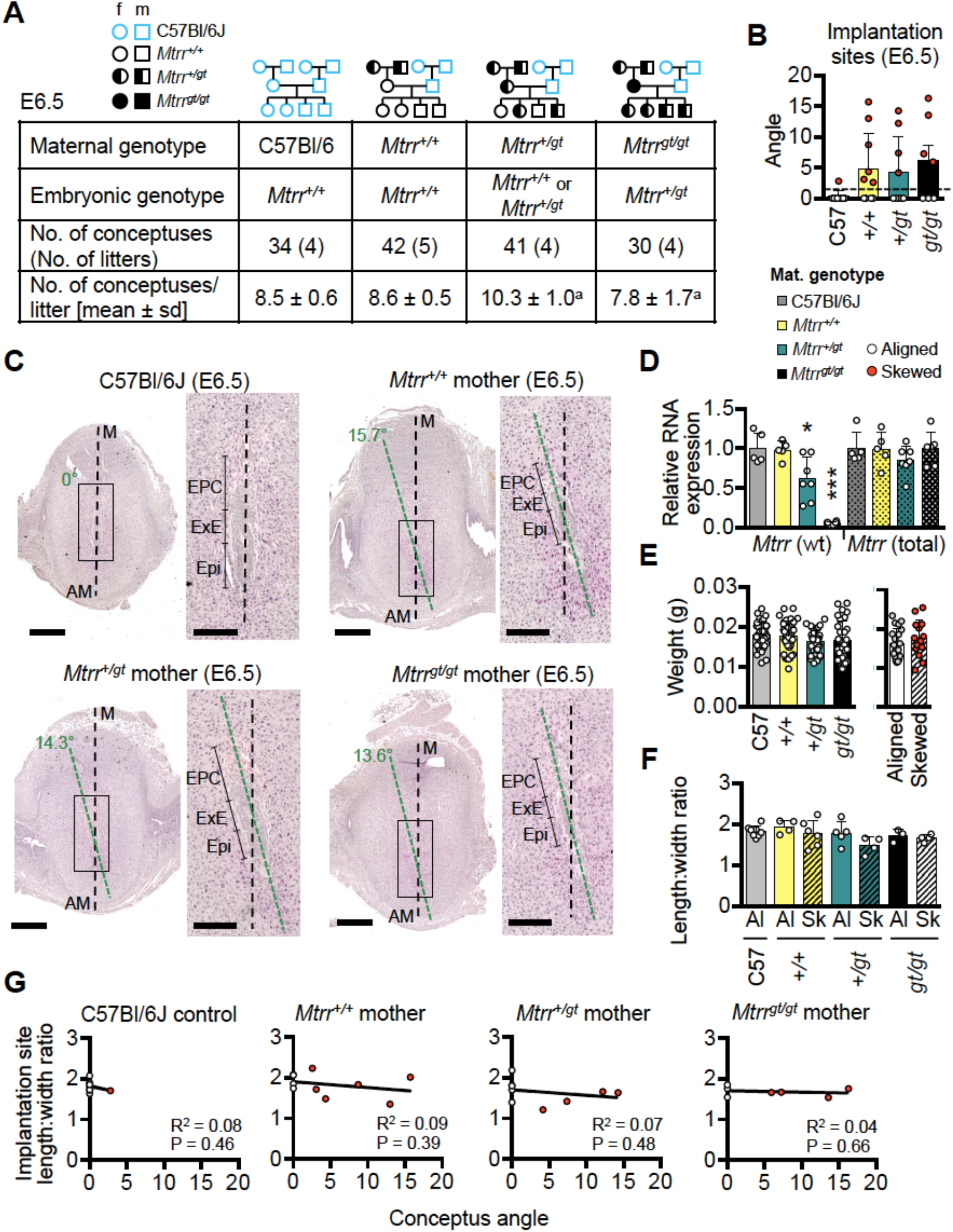
The effect of maternal *Mtrr*^*gt*^ genotype on mouse conceptus skewing at E6.5. (A) Litter parameters at E6.5 obtained when altering the maternal *Mtrr*^*gt*^ genotype. N=4 litters/cross with 31-41 conceptuses*/*cross. Litter sizes: Ordinary one-way ANOVA, P=0.0351, ^a^Dunn multiple comparison test: *Mtrr*^*+/gt*^ versus *Mtrr*^*gt/gt*^, P<0.05. Pedigree key: circle, female; squares, males; blue outline, C57Bl/6J mouse strain; black outline, *Mtrr*^*gt*^ mouse strain; white fill, *Mtrr*^*+/+*^; half black/half white, *Mtrr*^*+/gt*^; black fill, *Mtrr*^*gt/gt*^. (B) Graph showing the average angle of alignment at E6.5 in conceptuses derived from C57Bl/6J, *Mtrr*^*+/+*^, *Mtrr*^*+/gt*^, or *Mtrr*^*gt/gt*^ mothers and C57Bl/6J fathers. White dots, aligned conceptuses. Red dots, conceptuses with angles >2 sd above the control mean (indicated by dashed black line). N=7-10 conceptuses/group. Ordinary one-way ANOVA, P=0.1438. (C) H&E stained histological sections of implantation sites at E6.5 derived from C57Bl/6J, *Mtrr*^*+/+*^, *Mtrr*^*+/gt*^, or *Mtrr*^*gt/gt*^ mothers and C57Bl/6J fathers (N=7-10 conceptuses/group). Black dashed line bisects the implantation site. Green dashed line bisects conceptus. Angles indicate the degree of skewing. Boxed region indicates higher magnification image to the right-hand side. Scale bars, low magnification, 500 µm; high magnification, 250 µm. AM, antimesometrial pole; EPC, ectoplacental cone; Epi, epiblast; ExE, extra-embryonic ectoderm; M, mesometrial pole. (D) RT-qPCR analysis of relative RNA levels (mean ± sd) of the wildtype (wt) *Mtrr* transcripts and total *Mtrr* transcripts (wt + gene-trapped transcripts) in whole implantation sites at GD6.5 derived from C57Bl/6J, *Mtrr*^*+/+*^, *Mtrr*^*+/gt*^ and *Mtrr*^*gt/gt*^ mothers and C57Bl/6J fathers. N=5-7 implantation sites/group. Values were compared to C57Bl/6J (normalized to 1). One-way ANOVA with Tukey’s multiple comparison test, *P<0.05, ***P<0.001. (E) Left-hand graph: average weights of whole implantation sites at GD6.5 (mean ± sd) derived from C57Bl/6J, *Mtrr*^*+/+*^, *Mtrr*^*+/gt*^, or *Mtrr*^*gt/gt*^ mothers and C57Bl/6J fathers. N=30-42 conceptuses/group. Alignment not determined. One-way ANOVA, P=0.114. Right-hand graph: conceptus weights from *Mtrr*^*+/+*^, *Mtrr*^*+/gt*^, and *Mtrr*^*gt/gt*^ mothers were pooled and divided into aligned orientation (white dots, N=20 conceptuses) and skewed orientation (red dots; N=14 conceptuses). Independent t test, P=0.392. (F) Implantation site shape as determined by the ratio of decidua length (along the mesometrial-antimesometrial axis) and width (along the lateral axis). Maternal genotypes are indicated. Aligned (Al) and skewed (Sk) conceptuses were considered separately (N=3-8 implantation sites/group). Data presented as mean ± sd. One-way ANOVA, P=0.1371. (G) Linear regression analysis between conceptus angle and decidua length:width ratio for conceptuses at GD6.5 derived from C57Bl/6J, *Mtrr*^*+/+*^, *Mtrr*^*+/gt*^, and *Mtrr*^*gt/gt*^ mothers (N=7-10 conceptuses/group). White dots, aligned conceptuses; red dots, skewed conceptuses; dotted line, line of best fit.

### Gross decidual morphology is unchanged by maternal *Mtrr*^*gt*^ allele

Other studies showed that skewed conceptus orientation might result from abnormal decidual remodelling and signalling (Alexander et al., 1996; Zhang et al., 2014). Therefore, we assessed the effects of the maternal *Mtrr*^*gt*^ mutation on decidualisation. Since the extent of wildtype *Mtrr* mRNA knockdown in *Mtrr*^*+/gt*^ and *Mtrr*^*gt/gt*^ cells is tissue specific (Elmore et al., 2007; Padmanabhan et al., 2013; Sowton et al., 2020), wildtype *Mtrr* transcript levels were determined in whole implantation sites at gestational day (GD) 6.5 via RT-qPCR as a proxy for decidual cells (N=5-7 whole implantation sites/group). This was because implantation sites at GD6.5 consist largely of decidualized stroma with a relatively low contribution by fetal-derived cells. As expected, implantation sites from *Mtrr*^*+/+*^ mothers displayed a similar level of wildtype *Mtrr* transcripts as C57Bl/6J controls (98.0% of control levels; Figure 6D). Furthermore, this level was reduced to 62.0% and 6.0% of controls in implantation sites from *Mtrr*^*+/gt*^ and *Mtrr*^*gt/gt*^ mothers, respectively (P<0.0001; Figure 6D), and was within the range observed in other *Mtrr*^*+/gt*^ and *Mtrr*^*gt/gt*^ tissues (Elmore et al., 2007; Padmanabhan et al., 2013; Sowton et al., 2020). Total *Mtrr* transcript levels (included *Mtrr*^+^ and *Mtrr*^*gt*^ RNA) were similar across all maternal genotypes (P=0.4941; Figure 6D) indicating that transcriptional compensation for wildtype *Mtrr* deficiency did not occur.

Gross decidual morphology was assessed in histological sections of implantation sites at GD6.5. The extent of decidualisation was unaffected by a maternal *Mtrr*^*gt*^ allele at GD6.5 since the average implantation site weight did not differ between control and *Mtrr*^*+/+*^, *Mtrr*^*+/gt*^, or *Mtrr*^*gt/gt*^ mothers (Figure 6E; P=0.114). Furthermore, implantation site weight was similar between aligned and skewed conceptuses (Figure 6E; P=0.393), with no apparent correlation between implantation site weight and degree of conceptus skewing (Supplementary figures 1A,B). No litter size effect on implantation site weight at GD6.5 was observed (Supplementary figure 1C). Typically, an implantation site is ellipsoid with the mesometrial-antimesometrial axis (length) showing a longer dimension than the lateral axis (width). A rounder implantation site might indicate defective decidualisation (Alexander et al., 1996). Centrally-located transverse sections of implantation sites from C57Bl/6J mothers and from *Mtrr*^*+/+*^, *Mtrr*^*+/gt*^, or *Mtrr*^*gt/gt*^ mothers with aligned or skewed conceptuses (N=3-8 implantation sites/group) were measured along both axes. No significant differences in decidua length:width ratios were observed in implantation sites from any maternal genotypic group regardless of skewing (P=0.137; Figure 6F). Furthermore, there was no association between degree of conceptus skewing and implantation site dimensions (P>0.39; Figure 6G). When lateral decidual cell density was quantified by nuclear counts in implantation sites containing skewed conceptuses derived from *Mtrr*^*+/+*^, *Mtrr*^*+/gt*^, or *Mtrr*^*gt/gt*^ mothers (N=4-9 implantation sites/group), there was no significant difference compared to C57Bl/6J conceptuses (P=0.170; Supplementary figure 2C-D). Altogether, gross decidual morphology was unaffected by a maternal *Mtrr*^*gt*^ allele.

### Signalling pathways in decidua are potentially disrupted by the *Mtrr*^*gt*^ mutation

Next, we explored the effects of maternal *Mtrr*^*gt*^ allele on the molecular regulation of decidualization. Molecular markers of decidualisation were assessed by RT-qPCR in whole implantation sites at GD6.5 from C57Bl/6J, *Mtrr*^*+/+*^, *Mtrr*^*+/gt*^, and *Mtrr*^*gt/gt*^ mothers (N=5-7 sites/group). Progesterone receptor (PGR) (Lydon et al., 1995) and its downstream effectors NR2F2 (Wetendorf and DeMayo, 2012) and BMP2 (Lee et al., 2007) are indispensable for decidualisation. While *Pgr* and *Nr2f2* mRNA expression was significantly reduced in the implantation sites from *Mtrr*^*+/gt*^ and *Mtrr*^*gt/gt*^ mothers compared to C57Bl/6J and *Mtrr*^*+/+*^ mothers (P<0.0012; Figure 7A), downstream targets of PGR signalling including *Hand2* and *Hoxa10* (Lim et al., 1999; Okada et al., 2014) were unchanged (P=0.4132 and P=0.0796, respectively; Figure 7A). The degree of conceptus skewing was unknown in the samples assessed since each implantation site was snap frozen immediately after dissection, whereas analysis of conceptus alignment was determined in histological sections. However, immunohistochemistry suggested lower PGR protein levels in both *Mtrr*^*gt/gt*^ aligned and skewed conceptuses compared to C57Bl/6J controls (Figure 7B) though this decrease in expression might be insufficient to affect downstream effectors. Future studies should include protein quantification of PGR and downstream targets. Notably, *Bmp2* transcripts were significantly decreased in implantation sites from all three maternal genotypes relative to controls (P=0.0001; Figure 7A). Decreased *Bmp2* transcripts and conceptus skewing in *Mtrr*^*+/+*^ conceptuses derived from *Mtrr*^*+/+*^ mothers and *Mtrr*^*+/gt*^ maternal grandparents suggested that BMP2 signalling might have a role in conceptus alignment independent of progesterone signalling and decidualisation defects.

**Figure 7.**
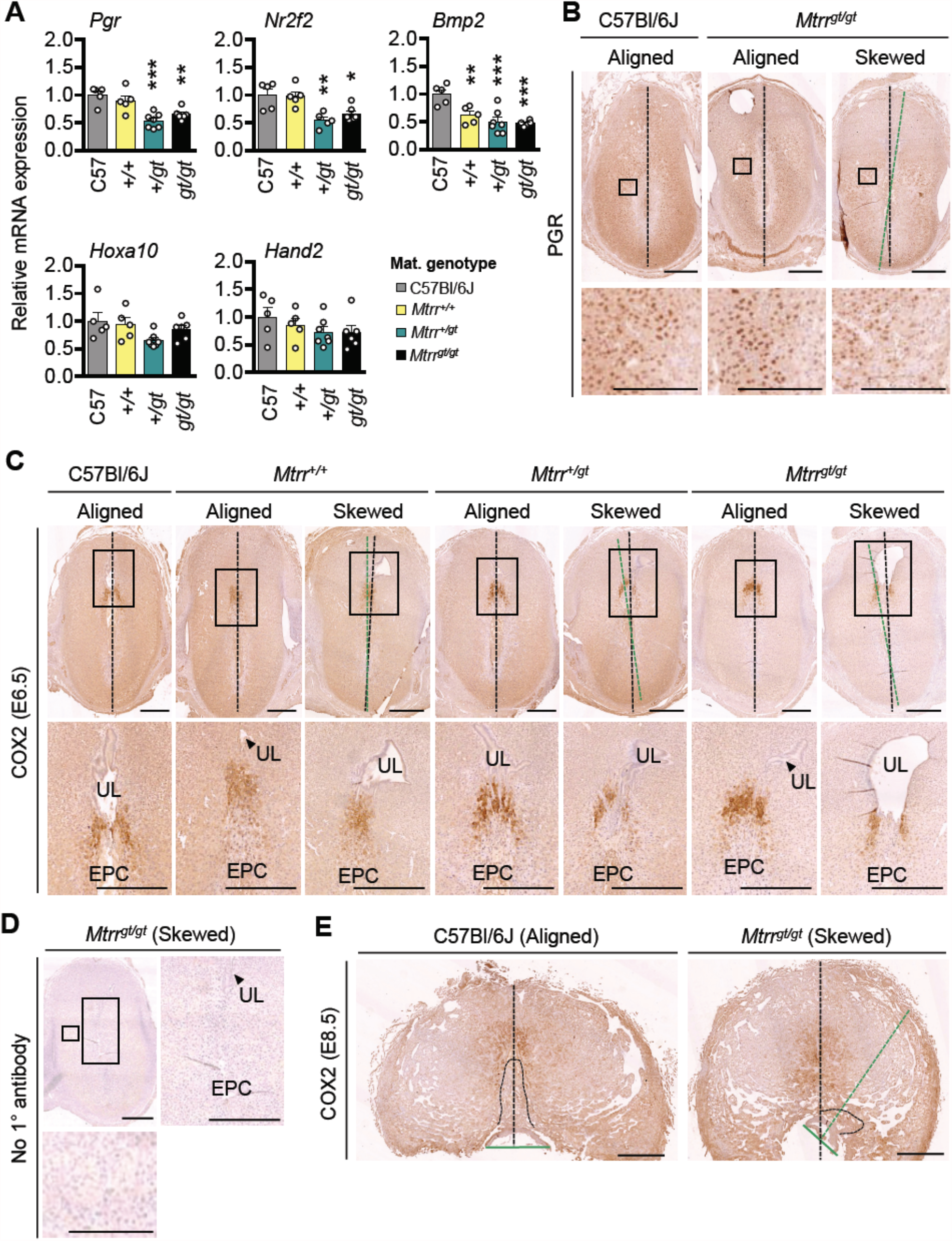
Maternal and grandparental *Mtrr*^*gt*^ allele might affect molecular signalling in mouse decidua at GD6.5. (A) RT-qPCR analysis of decidualisation gene markers in whole implantation sites at GD6.5 derived from C57Bl/6J, *Mtrr*^*+/+*^, *Mtrr*^*+/gt*^ or *Mtrr*^*gt/gt*^ mothers (N=5-7 sites/group). Data presented as mean ± sd, relative to C57Bl/6J controls (normalized to 1). One-way ANOVA with Tukey’s multiple comparison test, *P<0.05, **P<0.01, ***P<0.001. (B) PGR immunostaining in implantation sites at GD6.5 derived from C57Bl/6J mothers and aligned and skewed conceptuses from *Mtrr*^*gt/gt*^ mothers. (N=3 sites/genotype) Scale bars: 500 µm. (C) COX2 immunostaining (brown) in implantation sites at GD6.5 derived from C57Bl/6J, *Mtrr*^*+/+*^, *Mtrr*^*+/gt*^, or *Mtrr*^*gt/gt*^ mothers and C57Bl/6J fathers. N=2-5 sites/maternal genotype. DNA, blue. Maternal genotype is indicated. Both aligned and skewed conceptuses are shown. Scale bar: low magnification, 1 mm; high magnification, 500 µm. (D) No primary antibody control in C57Bl/6J implantation site at GD6.5. DNA, blue. Scale bar: low magnification, 1 mm; high magnification, (left) 500 µm; (bottom) 250 µm. (E) COX2 immunostaining (brown) in an aligned C57Bl/6J placenta and skewed *Mtrr*^*gt/gt*^ placenta at GD8.5. N=6 sites/genotype. DNA, blue. Dotted line, outline of EPC. Scale bar: 1 mm. (B-E) Boxed area indicates region of higher magnification. Black dotted line bisects the implantation site. Green dotted line bisects the conceptus (E6.5) or chorion (E8.5). EPC, ectoplacental cone. UL, uterine lumen remnant.

### Decidual COX2 expression is normal in implantation sites with skewed orientation

When the gene encoding for COX2 protein is knocked out, defects in implantation and decidualisation result (Lim et al., 1999). COX2 is important for prostaglandin synthesis and is initially expressed in the decidua adjacent to the implanting embryo at GD4.5 followed by restricted expression in the mesometrial decidua by GD5.5 and at GD7.5 (Chakraborty et al., 1996). Aberrant COX2 expression was also observed in other models of conceptus misorientation (Daikoku et al., 2011; Cha et al., 2014; Zhang et al., 2014). Therefore, we investigated COX2 protein expression in the decidua at E6.5 from *Mtrr*^*+/+*^, *Mtrr*^*+/gt*^, and *Mtrr*^*gt/gt*^ mothers (mated to C57Bl/6J males), and at E8.5 from *Mtrr*^*gt/gt*^ mothers (mated to *Mtrr*^*gt/gt*^ males), particularly in association with aligned and skewed conceptus orientation. Within centrally-located transverse histological sections, COX2 immunostaining in C57Bl/6J controls was restricted to the mesometrial decidua above the tip of the EPC, in line with the anitmesometrial-mesometrial axis (Figures 7C,E). A similar pattern was observed in implantation sites from *Mtrr*^*+/+*^, *Mtrr*^*+/gt*^ and *Mtrr*^*gt/gt*^ mothers, regardless of the extent of conceptus skewing (Figure 7C). Strikingly, the most skewed *Mtrr*^*gt/gt*^ conceptus at GD8.5 showed normal mesometrial decidual patterning of COX2 even though the EPC was more laterally located (Figure 7E). These data suggested that general decidual patterning was established and that COX2 expression did not dictate conceptus orientation in the *Mtrr*^*gt*^ model.

### Conceptus skewing is transgenerationally inherited in *Mtrr*^*gt*^ mouse line

The *Mtrr*^*gt*^ mouse line is a known model of TEI (Padmanabhan et al., 2013; Blake et al., 2021). Since conceptus skewing and low *Bmp2* transcript levels were apparent in *Mtrr*^*+/+*^ conceptuses at E6.5 derived from *Mtrr*^*+/+*^ parents and *Mtrr*^*+/gt*^ grandparents (Figures 6B,C, 7A), we explored a grandparental effect for misaligned orientation. To this end, we reanalysed our published data set at E10.5 whereby several genetic pedigrees were established to rigorously assess the effects of a maternal grandparental *Mtrr*^*gt*^ allele on the subsequent wildtype generations (Padmanabhan et al., 2013; Padmanabhan et al., 2017). Additional litters were included where possible. The breeding scheme was as follows: An *Mtrr*^*+/gt*^ female or male mouse (i.e., the F0 generation) was crossed with a control C57Bl/6J mate. Their wildtype daughters (i.e., the F1 generation) were then mated with a C57Bl/6J male and the wildtype grandprogeny (i.e., the F2 generation) were dissected and assessed at E10.5. Continuing a similar mating scheme allowed us to also assess the wildtype F3 and F4 generations. C57Bl/6J crosses were analysed as a control (see pedigree schematics in Figure 8).

**Figure 8.**
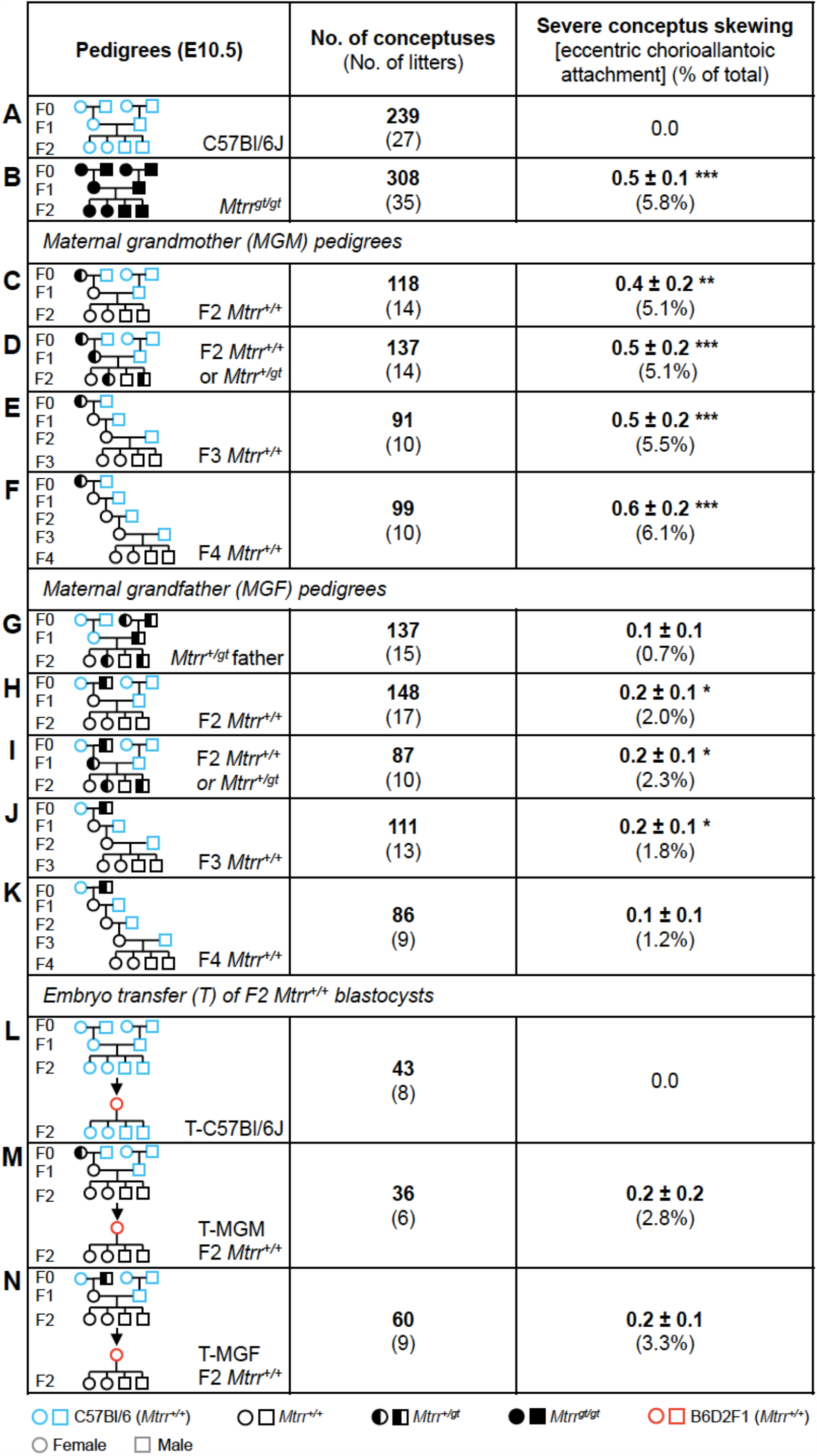
Severe conceptus skewing is transgenerationally inherited in the *Mtrr*^*gt*^ mouse line. (A-K) Frequency of eccentric chorioallantoic attachment at E10.5 as an indicator of severe conceptus skewing caused by the (C-F) *Mtrr*^*+/gt*^ maternal grandmother (MGM) or (H-K) *Mtrr*^*+/gt*^ maternal grandfather (MGF) compared to (A) C57Bl/6J conceptuses. (B) *Mtrr*^*gt/gt*^ and (G) *Mtrr*^*+/gt*^ paternal pedigrees were also assessed. (L-N) Frequency of eccentric chorioallantoic attachment at E10.5 resulting from the transfer of wildtype pre-implantation embryos derived from (M) an *Mtrr*^*+/gt*^ MGM and *Mtrr*^*+/+*^ mother (T-MGM) or (N) an *Mtrr*^*+/gt*^ MGF and *Mtrr*^*+/+*^ mother (T-MGF) into a B6D2F1 pseudopregnant recipient female. (L) C57Bl/6J embryos were transferred as a control. This experiment was originally performed in (Padmanabhan et al., 2013). Data is displayed as the average number of phenotypically affected conceptuses/litter (± sem) followed by the percentage of total conceptuses assessed in brackets. Independent t test compared to respective C57Bl/6J controls, *P<0.05, **P<0.01, ***P<0.001. Pedigree key: circle, female; squares, males; blue outline, C57Bl/6J mouse strain; red outline, B6D2F1 mouse strain; black outline, *Mtrr*^*gt*^ mouse strain; white fill, *Mtrr*^*+/+*^; black fill, *Mtrr*^*gt/gt*^; half black/half white, *Mtrr*^*+/gt*^.

The phenotypic frequency of ECA at E10.5 was used as an indicator of severe conceptus misalignment. Refer to our other published work for a full assessment of the phenotypic outcomes within the *Mtrr*^*gt*^ model (Padmanabhan et al., 2013; Padmanabhan et al., 2017). While we acknowledge that assessing only severe misalignment does not give a full picture of the extent of skewing in these pedigrees, the number of conceptuses assessed (N=86-308 conceptuses/pedigree) allowed for statistical analysis. Overall, a low frequency of ECA was observed in the wildtype F2 progeny of the *Mtrr*^*+/gt*^ maternal grandmother (MGM) pedigree (5.1% of conceptuses, P<0.01; Figure 8C) and the *Mtrr*^*+/gt*^ maternal grandfather (MGF) pedigree (2.0% of conceptuses, P<0.05; Figure 8H). This phenotype was not observed in the controls (Figure 8A). However, the *Mtrr* genotype of the F1 female in both the MGM and MGF pedigrees had no impact on the frequency of ECA. This is because the progeny of *Mtrr*^*+/+*^ or *Mtrr*^*+/gt*^ F1 females showed similar frequencies of ECA in their respective pedigrees (MGM: Figures 8C versus 8D; MGF: Figures 8H versus 8I). There was a negligible *paternal* effect of the *Mtrr*^*gt*^ allele on this phenotype (0.7% of conceptuses, P=0.183; Figure 8G). Notably, the frequency of ECA was greater in wildtype F2 progeny when derived from an *Mtrr*^*+/gt*^ MGM compared to an *Mtrr*^*+/gt*^ MGF (Figures 8C versus 8H) suggesting that the mechanism causing this phenotype might be different when initiated via the F0 oocyte versus the F0 sperm. The fact that *Mtrr*^*gt/gt*^ conceptuses at E10.5 derived from *Mtrr*^*gt/gt*^ intercrosses showed a similar frequency of ECA (5.8% of conceptuses, P<0.001) as wildtype F2 conceptuses in the MGM pedigree (Figures 8B versus 8C) indicated that the mechanism was similarly due to *Mtrr* deficiency in one (or both) maternal grandparent. Finally, we observed that ECA persisted until the wildtype F4 generation when derived from an *Mtrr*^*+/gt*^ F0 female (Figures 8E,F) yet only until the wildtype F3 generation when derived from an *Mtrr*^*+/gt*^ F0 male (Figures 8J,K). While the duration of phenotypic inheritance indicates TEI as a mechanism for conceptus misalignment in both pedigrees (Padmanabhan et al., 2013; Blake and Watson, 2016), phenotypic resolution occurred in the MGF pedigree. Whether phenotypic resolution occurs in the MGM pedigree after the F4 generation remains to be determined.

### Severe conceptus misalignment in the *Mtrr*^*gt*^ model likely occurs independent of the uterine environment

We previously reported that congenital malformations in wildtype F2 conceptuses derived from an *Mtrr*^*+/gt*^ MGM or MGF persisted after blastocysts were transferred into a normal F1 uterus (Padmanabhan et al., 2013). These blastocyst transfer experiments indicated that the cause of congenital malformations was independent of the F1 *Mtrr*^*+/+*^ uterine environment and likely the result of epigenetic inheritance via the germline (Padmanabhan et al., 2013). Re-analysis of this data showed that ECA (i.e., severe conceptus skewing) was among the phenotypes that persisted after blastocyst transfer (Figures 8M,N). The frequency did not reach statistical significance likely due to the low number of transferred embryos analysed (N=36-60 embryos from 6-9 litters). The control experiment, which involved the transfer of C57Bl/6J blastocysts into a normal uterus, did not exhibit ECA (Figure 8L) indicating that the blastocyst transfer protocol itself was not responsible for this phenotype. Overall, we propose that misaligned conceptus orientation occurred independent of a uterine environment effect. Instead, the mechanism was epigenetically inherited from a maternal grandparent and leading to a defect that was intrinsic to the embryo.

## DISCUSSION

It is well known that maternal folate deficiency is associated with increased risk for pregnancy complications. We have shown that abnormal folate metabolism due to the *Mtrr*^*gt*^ mutation in mice influences conceptus orientation and spacing within the uterus with implications for embryo and placenta development. These phenotypes are likely caused by a defect that is intrinsic to the embryo rather than the mother. Our data highlight the complexity of phenotype establishment associated with abnormal folate metabolism, with the mechanism acting beyond a direct maternal effect. Furthermore, we implicated the *Mtrr*^*gt*^ genotype of the maternal grandparents as the initial effector of conceptus misalignment. In general, the mechanism of TEI of phenotypes is not well understood and will take additional studies to elucidate. However, it is hypothesized that an abnormal epigenetic factor(s) is inherited via the germline to cause epigenetic instability in the offspring and to perpetuate phenotypic inheritance across multiple generations (see below; Padmanabhan et al., 2013; Blake and Watson, 2016; Blake et al., 2021).

We previously identified a phenotype in the *Mtrr*^*gt*^ mouse line whereby the allantois appeared to attach to the chorion at the wrong location when assessed via gross dissection and was originally characterised as ‘eccentric chorioallantoic attachment’ (Padmanabhan et al., 2013). Here, this phenotype is reclassified as ‘severe conceptus skewing’ meaning that the allantois attaches to the chorion in a normal manner, but the chorion and embryo/allantois are similarly ‘eccentrically’ oriented within the decidua. The establishment of mesometrial-antimesometrial conceptus orientation is not well understood. It might occur at implantation as the blastocyst attaches to the uterus and/or post-implantation as the developing conceptus invades into the decidualising endometrium and receives signals to direct its growth. These processes depend on extracellular matrix proteins, adhesion molecules, and cytokine signals (e.g., WNTs, FGFs, BMPs, etc.) (Red-Horse et al., 2004). Misalignment is most often associated with structural or molecular deficiencies in the uterus. For instance, aberrant formation of uterine crypts misorient the blastocyst at implantation (Daikoku et al., 2011; Cha et al., 2014; Zhang et al., 2014), decidualisation defects associate with misalignment of conceptuses post-implantation (Alexander et al., 1996; Zhang et al., 2014), and disorganisation of the decidual vasculature network and increased blood perfusion misdirects trophoblast invasion resulting in skewed growth (Winterhager et al., 2013). The establishment of signalling gradients in decidua (e.g., WNT5) are likely also required for conceptus alignment (Daikoku et al., 2011; Cha et al., 2014; Goad et al., 2017). Evidence presented here argues against a uterine defect as the main cause of conceptus misalignment in the *Mtrr*^*gt*^ model since normal morphology and patterning of *Mtrr*^*gt/gt*^ decidua was observed, and skewed *Mtrr* conceptuses were present after blastocyst transfer into a normal uterus. While down-regulation of *Bmp2* mRNA in the decidua of all *Mtrr* maternal genotypes at E6.5 was evident, further experiments in the *Mtrr*^*gt*^ model are required to explore whether this and other uterine signalling pathways directly influence conceptus orientation from implantation onwards.

Alternatively, an embryo-specific defect is more likely to cause conceptus misalignment in the *Mtrr*^*gt*^ model. Mouse blastocysts attach to the uterine epithelium via adhesion molecules (e.g., integrins, L-selectins) expressed on the mural trophectoderm (Cross et al., 1994; Red-Horse et al., 2004) so that the embryonic-abembryonic axis is parallel to the mesometrial-antimesometrial axis (Smith, 1985). How the blastocyst attaches to the uterine epithelium under normal circumstances is not well understood. It is hypothesized that the blastocyst might attach in any orientation and then roll along the uterine epithelium into the correct orientation (Kirby et al., 1967; Red-Horse et al., 2004). Therefore, it is possible that critical adhesion molecules on the mural trophectoderm of *Mtrr* blastocysts are regionally expanded or misexpressed resulting in skewed blastocyst attachment to the uterine epithelium. Given normal litter sizes from *Mtrr*^*gt/gt*^ intercrosses are apparent at midgestation (this study; Padmanabhan et al., 2013), implantation of *Mtrr*^*gt/gt*^ blastocysts can occur regardless of the proposed adhesion defect. Alternatively, after normal blastocyst orientation at implantation, some conceptuses from the *Mtrr*^*gt*^ mouse line might be unresponsive to cytokine signals from the decidua that guide mesometrial-antimesometrial alignment. Further studies are required to elucidate the specific embryocentric mechanisms and molecular pathways involved in conceptus orientation, and how they are influenced by abnormal folate metabolism.

It is probable that conceptus misalignment negatively impacts embryo and placenta development. The degree of skewing occurred over a continuum, though whether minor skewing is harmful to development is currently unclear. Even though normal trophoblast lineage patterning (Simmons et al., 2008) occurred in severely skewed *Mtrr*^*gt/gt*^ conceptuses at E8.5, an improperly oriented placenta would likely have insufficient access to maternal blood for normal fetoplacental development. Indeed, we observed that severe conceptus skewing at E10.5 was associated with embryonic developmental delay and heart malformations. Others have proposed the placenta-heart axis whereby that primary placental phenotype can result in secondary cardiovascular abnormalities (Barak et al., 1999; Hemberger and Cross, 2001; Perez-Garcia et al., 2018), and vice versa (Outhwaite et al., 2019). Whether this phenomenon occurs in skewed conceptuses is unclear and requires further investigation by conditional mutation or tetraploid aggregation experiments. Notably, neural tube closure defects, which are famously associated with maternal folate deficiency, did not correlate with conceptus misalignment indicating a separate causative mechanism.

Severe conceptus misorientation in the *Mtrr*^*gt*^ mouse line may show some relationship to velamentous umbilical cord insertion in humans. This condition is when the umbilical cord is misaligned and inserts into the chorioamniotic membranes at the edge of the placenta (Jauniaux et al., 2020) leading to reduced vascularisation of the placenta (Yampolsky et al., 2009). Its occurrence is associated with congenital heart disease and adverse perinatal outcomes including preterm birth and low birth weight (Raisanen et al., 2012; Ebbing et al., 2013; Brouillet et al., 2014; Albalawi et al., 2017). Importantly, the site of cord insertion might be determined by blastocyst orientation at implantation (Ismail et al., 2017), which we have proposed as a possible mechanism for conceptus misalignment in the *Mtrr*^*gt*^ model. Further study is required to elucidate similarities between these two conditions, particularly in the context of abnormal folate uptake or metabolism.

Dizygotic twinning is a seemingly rare event in mice. It occurs independent of the number of ovulations and likely results from poor blastocyst spacing at implantation. Indeed, excessive intrauterine fluid disrupts embryo spacing and infrequently leads to twinning due to abnormal interaction between the uterine epithelium and blastocysts (Lu et al., 2013). Typically, blastocyst spacing in the mouse uterus is a highly regulated event requiring mechanical processes (e.g., muscular contractions, ciliary movement) and molecular signalling between the uterus and blastocyst (Shi et al., 2014; Flores et al., 2020). Also, inhibitory signalling between neighbouring pre-implantation littermates might transpire. The molecular mechanisms of blastocyst spacing are not well understood. In general, normal spacing of *Mtrr*^*gt/gt*^ conceptuses along the length of the *Mtrr*^*gt/gt*^ uterus was apparent (this study), potentially ruling out a mechanical defect in the uterus (Flores et al., 2020). Indeed, the low frequency of twinning in the *Mtrr*^*gt*^ mouse line implies an embryo-specific mechanism since a maternal effect would affect a greater number of conceptuses within one litter. It is unclear whether maternal folate deficiency or abnormal folate metabolism resulting from a genetic mutation cause a similar incidence of twinning, though it is possible that this phenotype is unique to the *Mtrr*^*gt*^ mouse line. Although, other models of conceptus misalignment display extensive embryo spacing defects (Daikoku et al., 2011; Cha et al., 2014), it is unclear whether poor responsiveness of the blastocyst to maternal signals is a common mechanism between twinning and orientation phenotypes in the *Mtrr*^*gt*^ model.

Uterine structure at implantation and the frequency of blastocyst implantation are unaffected by dietary folate deficiency in mice (Gao et al., 2012), though post-implantation decidualisation and deciduoma formation might be inhibited (Geng et al., 2015). Since decidualisation was unaffected by the *Mtrr*^*gt/gt*^ mutation at E6.5, the effects of abnormal folate metabolism caused by the *Mtrr*^*gt*^ allele and dietary folate deficiency might differ. Conceptus misalignment and twinning are yet-to-be reported in other genetic mouse models whereby folate uptake or metabolism is disrupted. However, several of these models undergo embryonic lethality in early to mid-gestation (Piedrahita et al., 1999; Swanson et al., 2001; Gelineau-van Waes et al., 2008; Pickell et al., 2009; Salojin et al., 2011) suggesting that conceptus orientation should be explored in these models. Better understanding of the role of folate metabolism in this process is required to assess its implications for assisted reproductive technologies and early pregnancy loss.

One-carbon metabolism is involved in thymidine synthesis and cellular methylation reactions (Ducker and Rabinowitz, 2017). Thymidine synthesis is likely unaffected by the *Mtrr*^*gt*^ mutation since *de novo* mutations, which transpire from low thymidine levels resulting in uracil misincorporation into DNA (Blount et al., 1997), occur at a similar frequency in C57Bl/6J control and *Mtrr*^*gt/gt*^ mice (Blake et al., 2021). However, considerable alteration in patterns of DNA methylation was identified in adult germ cells and somatic tissues, and in placentas of the *Mtrr*^*gt*^ model, with some methylation changes associated with gene misexpression (Padmanabhan et al., 2013; Bertozzi et al., 2021; Blake et al., 2021). The broad range of developmental phenotypes observed in the *Mtrr*^*gt*^ model suggests that this epigenetic instability occurs stochastically on an inter-individual basis, ultimately leading to disruption of different gene pathways and different phenotypes. This stochasticity might explain why only some conceptuses are misaligned. It also creates a challenge when identifying specific genomic regions and target genes affected by abnormal folate metabolism that might transcriptomically explain the phenotypes that arise.

Importantly, severe conceptus skewing at E10.5 in the *Mtrr*^*gt*^ model exhibits a pattern of transgenerational inheritance. This non-conventional mode of inheritance is caused by exposure to an environmental stressor or metabolic disruption only in the initiating F0 generation and occurs independent of DNA base-sequence mutations (Blake and Watson, 2016). The *Mtrr*^*gt*^ mouse line is a known model of TEI as several embryonic phenotypes are inherited by the wildtype grandprogeny and great grandprogeny when either maternal grandparent is a carrier for the *Mtrr*^*gt*^ mutation (Padmanabhan et al., 2013; Blake et al., 2021). While the mechanism of epigenetic inheritance of phenotypes is unknown in the *Mtrr*^*gt*^ model and other mammalian multigenerational models, it is possible that epigenetic disruption in germ cells leads to epigenetic instability and transcriptional defects in the generations that follow (Blake and Watson, 2016). DNA methylation, histone modifications, and small non-coding RNA misexpression are proposed epigenetic candidates (Chen et al., 2016; Sharma et al., 2016; Blake et al., 2021; Lismer et al., 2021), but further experiments are required to elucidate the specific mechanism of TEI in the *Mtrr*^*gt*^ model. We discerned a greater frequency and later resolution of conceptus misalignment in wildtype conceptuses at E10.5 derived from an *Mtrr*^*+/gt*^ maternal grandmother versus an *Mtrr*^*+/gt*^ maternal grandfather. These observations might hold clues for future mechanistic exploration.

The association between folate metabolism and conceptus alignment has implications for early pregnancy loss associated with folate deficiency in humans, and for culture conditions involved in assisted reproductive technologies. Since conceptus skewing occurred several wildtype generations after the initial metabolic disruption, it is possible that unexplained pregnancy loss in humans might be associated with folate deficiency in the grandparental generation. Our study highlights the importance of folate in healthy pregnancies, yet emphasises that its role is more complex than a direct maternal effect.

## AUTHOR CONTRIBUTION

EDW: conceptualization, experimental design, and funding acquisition. KM, JR, and EDW: dissections, phenotyping, and histology preparation. KM: RNA isolation and RT-qPCR. ALW, KM, JR, and XST: histological staining, immunostaining and in situ hybridization. ALW, KM, and EDW: data analysis. ALW, EDW: data interpretation and manuscript writing. ALW, EDW, KM: manuscript editing. All authors contributed to the article and approved the submitted version.

## ACKNOWLEDGEMENTS

We thank Dr Simon Tunster for technical advice and support, and Dr Claire Senner and Prof Kathy Niakan for critical discussion of the manuscript.

## SUPPLEMENTARY MATERIALS

### SUPPLEMENTARY TABLES

**Supplementary Table 1.**
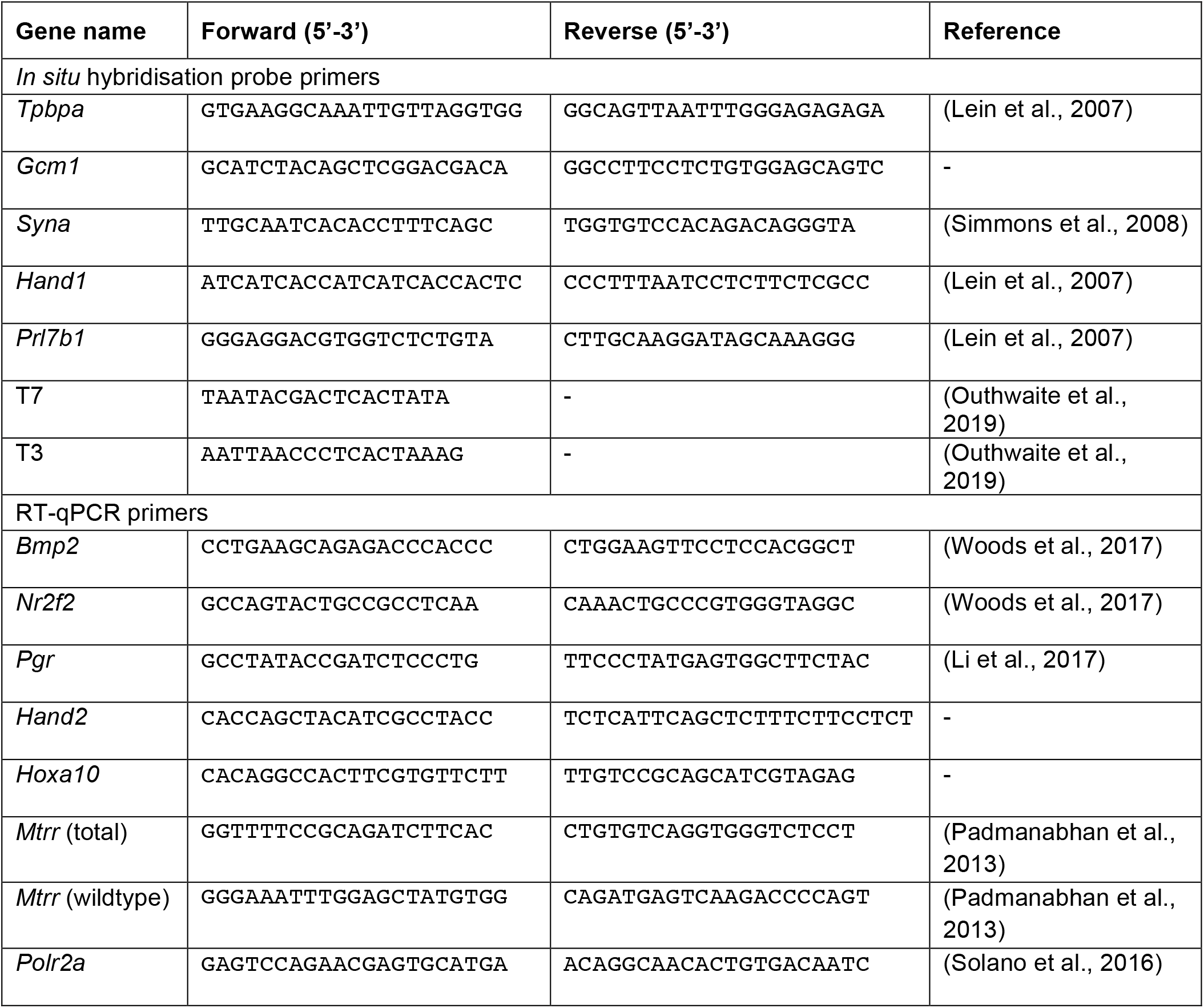
*In situ* hybridisation probe primers and RT-qPCR primers

### SUPPLEMENTARY FIGURES

**Supplementary figure 1.**
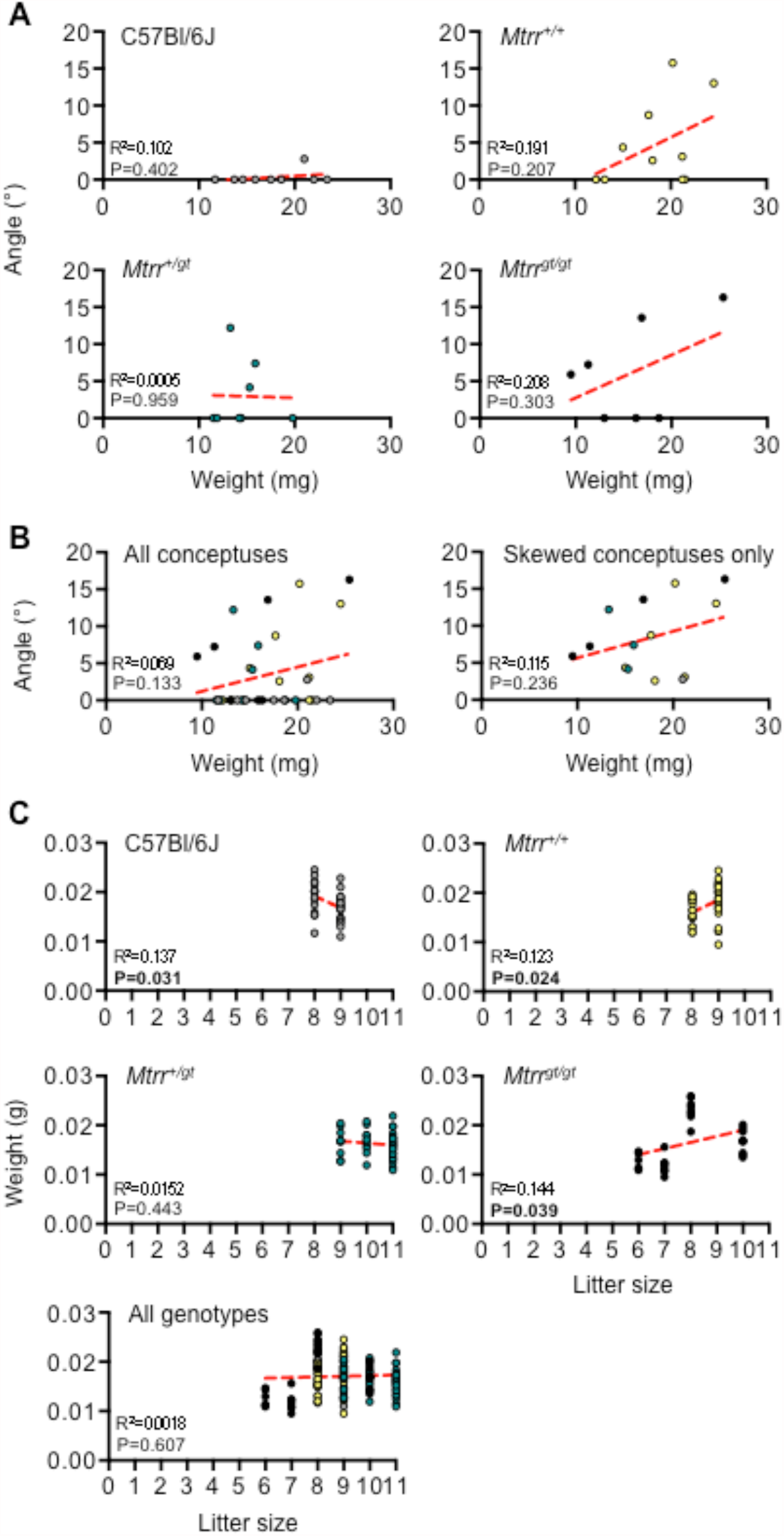
No correlation between *Mtrr*^*gt/gt*^ implantation site weight and degree of skewing or litter size at GD6.5. (A-B) Linear regression analyses between whole implantation site weight and angle of conceptus relative to the central midline. (A) Weights and angles were measured in conceptuses derived from C57Bl/6J (grey dots), *Mtrr*^*+/+*^ (yellow dots), *Mtrr*^*+/gt*^ (green dots), and *Mtrr*^*gt/gt*^ (black dots) females mated with C57Bl/6J males. The maternal genotype is shown. (B) The data from (A) was pooled together (all conceptuses, left-hand graph) to increase sample size. Similarly, the data from the misaligned conceptuses (>2 sd above the mean conceptus angle in C57Bl/6J conceptuses, see Fig. 5B) were extracted and plotted to determine whether there was a correlation between implantation site weight and degree of misalignment. Red line indicates line of best fit. (C) Linear regression analyses of litter size and implantation site weight at GD6.5 in conceptuses from C57Bl/6J (grey dots), *Mtrr*^*+/+*^ (yellow dots), *Mtrr*^*+/gt*^ (green dots), and *Mtrr*^*gt/gt*^ (black dots) females mated with C57Bl/6J males. The maternal genotype is shown. Red line indicates line of best fit.

**Supplementary figure 2.**
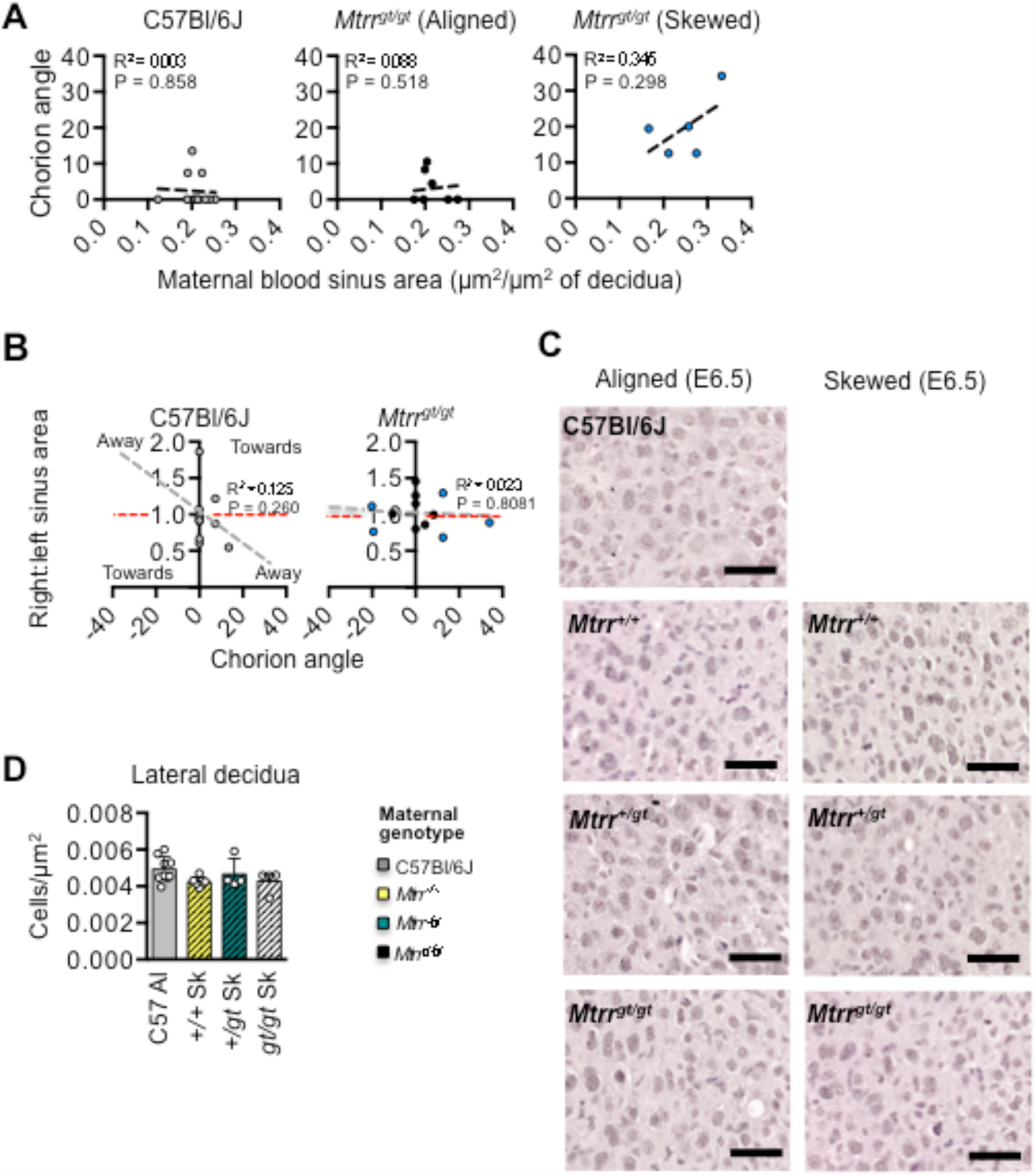
Normal decidual blood spaces, decidual cell morphology and density regardless of *Mtrr*^*gt*^ maternal genotype. (A) Linear regression analyses of decidua blood sinus area in relation to chorion angle for C57Bl/6J (white circles) and *Mtrr*^*gt/gt*^ conceptuses at GD8.5. Aligned (black circles) and skewed (blue circles) *Mtrr*^*gt/gt*^ placentas were considered separately (N=6-12 conceptuses/group). Dotted line indicates line of best fit. See also Figure 5E. (B) Linear regression analyses between the chorion angle and the ratio of right:left decidua blood sinus area in C57Bl/6J and *Mtrr*^*gt/gt*^ conceptuses at E8.5 (N=6-12 conceptuses/group). Aligned (grey or black dots) and skewed (blue dots) conceptuses are shown. Dashed red line, right:left blood sinus area ratio of 1. Dashed grey line, line of best fit. See also Figure 5G. (C) Histological sections stained with H&E showing lateral decidua in implantation sites at GD6.5 derived from C57Bl/6, *Mtrr*^*+/+*^, *Mtrr*^*+/gt*^, and *Mtrr*^*gt/gt*^ females and C57Bl/6J males. The maternal genotype is indicated. Decidua associated with conceptuses that were aligned to or skewed from the midline is shown. Scale bars: 50 µm. (D) Graph showing the average number of lateral decidual cells per µm^2^ in histological sections from C57Bl/6 (grey bar), *Mtrr*^*+/+*^ (yellow bar), *Mtrr*^*+/gt*^ (green bar), and *Mtrr*^*gt/gt*^ (black bar) females mated with C57Bl/6J males. Data is shown as mean ± sd. N=4-8 implantation sites/group with at least three histological sections assessed per individual. Al, aligned; Sk, skewed. One-way ANOVA.

